# Physical and functional convergence of the autism risk genes *Scn2a* and *Ank2* in neocortical pyramidal cell dendrites

**DOI:** 10.1101/2022.05.31.494205

**Authors:** Andrew D. Nelson, Amanda M. Catalfio, Julie M. Gupta, Lia Min, Rene N. Caballero-Floran, Kendall P. Dean, Carina C. Elvira, Kimberly D. Derderian, Henry Kyoung, Atehsa Sahagun, Stephan J. Sanders, Kevin J. Bender, Paul M. Jenkins

## Abstract

Dysfunction in sodium channels and their ankyrin scaffolding partners have both been implicated in neurodevelopmental disorders, including autism spectrum disorder (ASD). In particular, the genes SCN2A, which encodes the sodium channel NaV1.2, and ANK2, which encodes ankyrin-B, have strong ASD association. Recent studies indicate that ASD-associated haploinsufficiency in Scn2a impairs dendritic excitability and synaptic function in neocortical pyramidal cells, but how NaV1.2 is anchored within dendritic regions is unknown. Here, we show that ankyrin-B is essential for scaffolding NaV1.2 to the dendritic membrane of mouse neocortical neurons, and that haploinsufficiency of Ank2 phenocopies intrinsic dendritic excitability and synaptic deficits observed in Scn2a+/- conditions. Thus, these results establish a direct, convergent link between two major ASD risk genes and reinforce an emerging framework suggesting that neocortical pyramidal cell dendritic dysfunction can be etiological to neurodevelopmental disorder pathophysiology.

## INTRODUCTION

A decade of gene discovery has identified hundreds of genes whose dysfunction is associated with autism spectrum disorder (ASD) (Iossifov et al., 2014; Neale et al., 2012; O’Roak et al., 2012; Sanders et al., 2015, 2012; Satterstrom et al., 2020). A key challenge remains to translate these findings into an understanding of pathophysiology at the cellular and circuit level. Loss-of-function in *SCN2A*, which encodes the neuronal sodium channel NaV1.2, has the strongest evidence of ASD association based on exome sequencing (Ben-Shalom et al., 2017; Fu et al., 2021; Satterstrom et al., 2020; Spratt et al., 2019). Given this critical role in ASD etiology, alterations in cellular function due to *SCN2A* loss may illuminate common causes of dysfunction shared with other ASD-associated genes.

A novel role for Na_V_1.2 was identified recently in neocortical pyramidal cells, a cell class whose dysfunction is implicated in ASD (Satterstrom et al., 2020; Willsey et al., 2013). In contrast to the well-characterized roles for sodium channels (Na_V_s) in axonal action potential (AP) electrogenesis and propagation, Na_V_1.2 was found to be critical for dendritic excitability, with ASD-associated *Scn2a* haploinsufficiency impairing postsynaptic features of synaptic function and plasticity (Spratt et al., 2019). This dendritic Na_V_1.2 localization is presumably controlled by ankyrins, which are a family of intracellular scaffolding proteins that link ion channels to the underlying actin cytoskeleton (Bennett and Lorenzo, 2016; Lemaillet et al., 2003; Nelson and Jenkins, 2017). Ankyrin-Na_V_ interactions have been studied extensively in excitable axonal compartments, where Na_V_s are anchored by ankyrin-G (*ANK3*) (Jenkins and Bennett, 2001; Jenkins et al., 2015; Pan et al., 2006; Zhou et al., 1998), but how Na_V_s are scaffolded to dendritic domains to regulate postsynaptic excitability is unknown.

Insight into this question may come from ASD gene discovery, where another ankyrin family member, ankyrin-B (*ANK2*), has strong evidence of ASD association (Fu et al., 2021; Satterstrom et al., 2020). Interestingly, immunostaining for ankyrin-G and ankyrin-B in cultured neurons indicates that they occupy largely non-overlapping domains, with ankyrin-G enriched in the axon initial segment (AIS) and nodes of Ranvier, and ankyrin-B enriched in other regions, including dendrites (Lorenzo et al., 2014). Thus, ankyrin-B is well-positioned to scaffold dendritic Na_V_1.2 channels. In this way, loss-of-function in either *SCN2A* or *ANK2* could impair dendritic excitability, either directly through reduced Na_V_ density or function or indirectly by reduced Na_V_ scaffolding, respectively. This would implicate dendritic excitability as a convergent feature disrupted in ASD.

Here, we paired cellular and molecular biology with electrophysiology and two-photon imaging to demonstrate that the protein products of these two ASD risk genes, *SCN2A* and *ANK2*, interact in neocortical pyramidal cell dendrites to mutually regulate dendritic excitability. Using a novel epitope-tagged Na_V_1.2, we found that Na_V_1.2 co-localizes with ankyrin-B in the dendrites of mature neocortical neurons. Removal of ankyrin-B eliminated Na_V_1.2 dendritic localization. Furthermore, dendritic ankyrin-B loss was not compensated for by other ankyrin family members, indicating that ankyrin-B has a unique scaffolding role in this neuronal compartment. *Ex vivo* studies revealed that *Ank2* haploinsufficiency results in intrinsic and synaptic dendritic deficits that closely phenocopy those observed in *Scn2a* heterozygous neurons. Thus, these findings suggest that deficits in dendritic excitability may be a common point of convergence in ASD, with direct convergence between two high-risk genes *SCN2A* and *ANK2*.

## RESULTS

### Na_V_1.2 and ankyrin-B colocalize in mature neocortical neuron dendrites

To determine whether Na_V_s and ankyrins physically converge in dendrites, we first aimed to visualize their subcellular distribution patterns within a single cell. While Na_V_s can be observed in the AIS and nodes of Ranvier using conventional immunostaining approaches, they are often too diffuse to detect reliably in other regions (Inda et al., 2006). To visualize the subcellular localization of Na_V_1.2 across development in all neuronal compartments, we first generated a cDNA encoding full-length Na_V_1.2 with a 3xFLAG epitope tag on the carboxyl terminal, in addition to IRES-mediated expression of freely diffusible GFP. This Na_V_1.2-3xFLAG-IRES-GFP cDNA was created by cloning a codon-optimized *SCN2A* cDNA previously used to characterize ASD-associated variants (Ben-Shalom et al., 2017). Voltage-clamp recordings from HEK293 cells transfected with either wild type (WT) Na_V_1.2 or Na_V_1.2-3xFLAG indicated that the introduction of this epitope tag did not alter channel biophysics (**Figure S1A**) or its ability to interact with β1 subunits (**Figure S1B**), suggesting that this approach is a viable way to visualize Na_V_1.2 without altering its function.

Immunostaining and electrophysiological measurements indicate that Na_V_1.2 is enriched in the AIS of neocortical pyramidal cells in early development (Boiko et al., 2003; Gazina et al., 2015; O’Brien and Meisler, 2013; van Wart et al., 2007) (**Figure 1A**). Consistent with staining of native channels, Na_V_1.2-3xFLAG was similarly restricted to the AIS at day in vitro 7 (DIV7) in cultured neocortical neurons, where it colocalized with ankyrin-G (**Figure 1C**). Later in development, Na_V_1.2 is largely displaced from the AIS and instead increases in density throughout somatodendritic domains (Hu et al., 2009; Lorincz and Nusser, 2010; Spratt et al., 2019, 2021; Zhang et al., 2021) (**Figure 1A**). Consistent with this shift in Na_V_1.2 subcellular localization, Na_V_1.2-3xFLAG was visualized at high levels throughout dendrites at DIV21 (**Figure 1C**). Dendritic Na_V_1.2 was not co-localized with ankyrin-G (*ANK3)* (**Figure 1C**), suggesting that another ankyrin may be important for Na_V_ scaffolding in dendritic regions. Based on genetic and co-expression data related to ASD (Fu et al., 2021; Satterstrom et al., 2020; Willsey et al., 2013) (**Figure 1B**), we hypothesized that this dendritic scaffold is ankyrin-B (*ANK2)*. To compare the subcellular localization patterns of ankyrin-B and Na_V_1.2 across development, we transfected cultured neocortical neurons with Na_V_1.2-3xFLAG-IRES-GFP and immunostained with antibodies against ankyrin-B at DIV7 and DIV21. At DIV7, ankyrin-B was predominantly localized to the distal axon (**Figure 1D**). At DIV21, however, ankyrin-B was enriched along the membrane of dendritic shafts, colocalizing with Na_V_1.2 (**Figure 1D**). Importantly, there was no correlation between dendritic Na_V_1.2-3xFLAG and plasmid expression levels, inferred from GFP fluorescence intensity, suggesting that Na_V_1.2 expression is tightly regulated and that its dendritic localization is not an off-target effect of overexpression (**Figure 1E**). These data indicate ankyrin-B is well-positioned to scaffold Na_V_1.2 to the dendritic membrane of mature neocortical pyramidal neurons.

**Figure 1:**
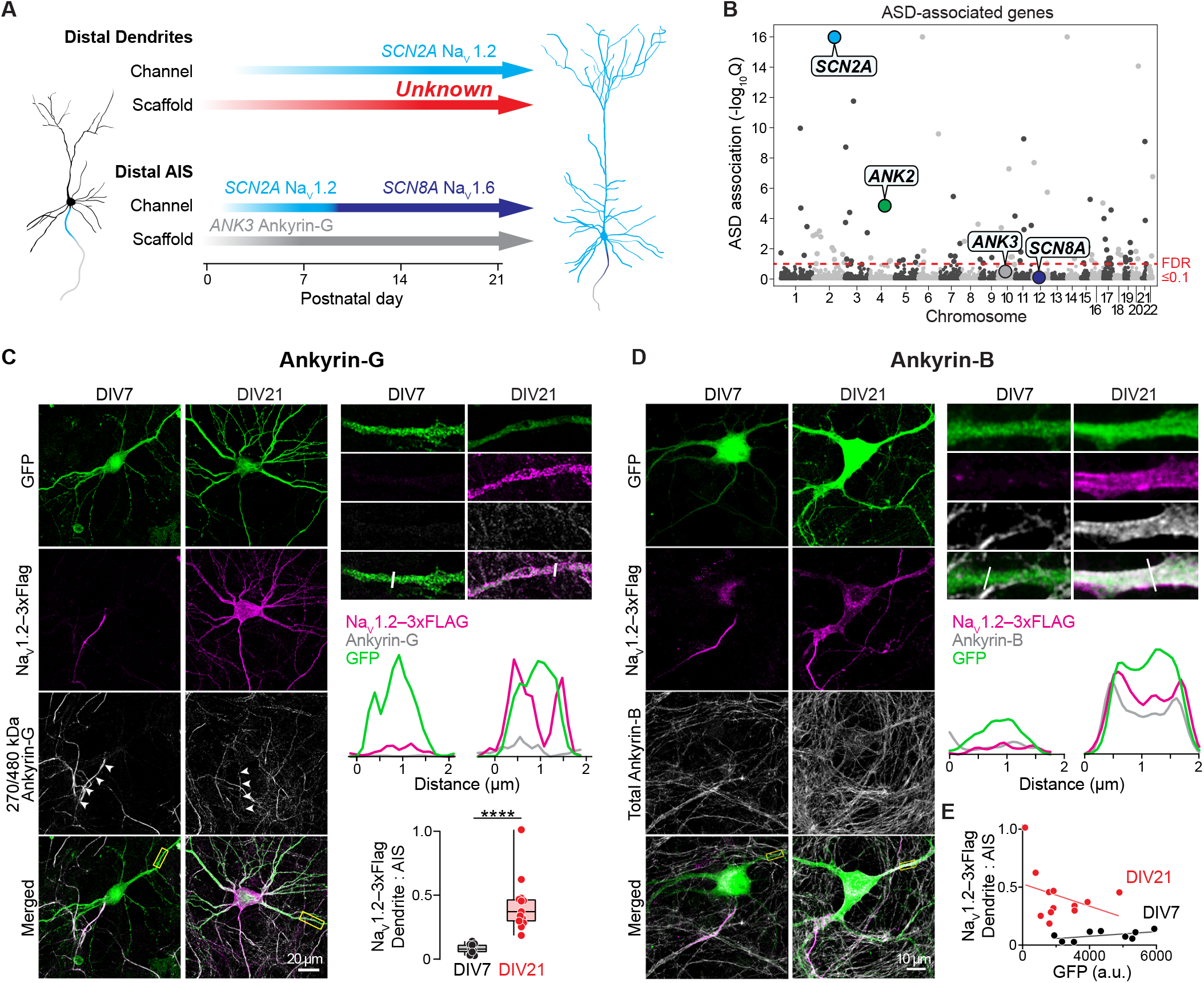
Ankyrin-B is colocalized with Na_V_1.2 in the dendrites of mature pyramidal neurons. **A**. Schematic of NaV channel distribution with ankyrin scaffolds throughout neocortical pyramidal neuron development. **B**. Manhattan plot of ASD-associated genes identified by whole-exome sequencing data. Red dotted-line indicated false discovery rate (FDR) threshold of ≤ 0.1. **C**. Left: Confocal images of wild-type cultured neocortical neurons transfected with NaV1.2-3xFLAG-IRES-eGFP and fixed at 7DIV or 21DIV. Cells were immunostained with antibodies against ankyrin-G (white), FLAG (magenta), and GFP (green). White arrows denote AIS of transfected cell. Right: zoomed image of dendrite labeled with yellow box. Plot profiles of dendritic NaV1.2-3xFLAG, ankyrin-G, and GFP fluorescence intensity (a.u.) at 7DIV (left) and 21DIV (right) determined by white line above. Quantification of mean fluorescence intensity of NaV1.2-3xFLAG in dendrites relative to AIS. Circles represent individual neurons. (7DIV: 0.08 ± 0.01, n = 10 cells; 21DIV: 0.4 ± 0.06, n = 13 cells) ****p < 0.0001. Mann-Whitney test. **D**. Left: Confocal images of cultured neocortical neurons transfected with NaV1.2-3xFLAG-IRES-eGFP and fixed at 7DIV or 21DIV. Cells were immunostained with antibodies against total ankyrin-B (white), FLAG (magenta), and GFP (green). Right: zoomed image of dendrite labeled with yellow box. Plot profiles of dendritic NaV1.2-3xFLAG, ankyrin-B, and GFP fluorescence intensity (a.u.) at 7DIV (left) and 21DIV (right) determined by white line above. **E**. Linear regression of mean NaV1.2-3xFLAG dendrite:AIS fluorescent signal versus GFP at DIV7 and DIV21. Note that dendritic localization of NaV1.2 occurs across a wide range of plasmid expression.

### Ankyrin-B localizes Na_V_1.2 to the dendritic membrane of mature neocortical neurons

To determine whether ankyrin-B directly scaffolds Na_V_1.2 to the dendritic membrane, we performed knockout-and-rescue experiments in cultured neocortical neurons generated from *Ank2*^*fl/fl*^ mice, which contain loxP sites flanking exon 24 of the *Ank2* gene (Roberts et al., 2019). In wild type (WT) conditions, Na_V_1.2-3xFLAG was highly enriched in the dendritic membrane with endogenous ankyrin-B in DIV21 neurons (**Figure 2A**). Co-transfection of Na_V_1.2-3xFLAG-IRES-GFP with Cre-2A-BFP (blue fluorescent protein) in *Ank2*^*fl/fl*^ neurons resulted in the complete absence of ankyrin-B and a corresponding loss of dendritic Na_V_1.2 (**Figure 2A**). Simultaneous knockout of endogenous ankyrin-B via Cre-2A-BFP and rescue with the canonical wild-type ankyrin-B restored Na_V_1.2 to the dendritic membrane (**Figure 2A**).

**Figure 2:**
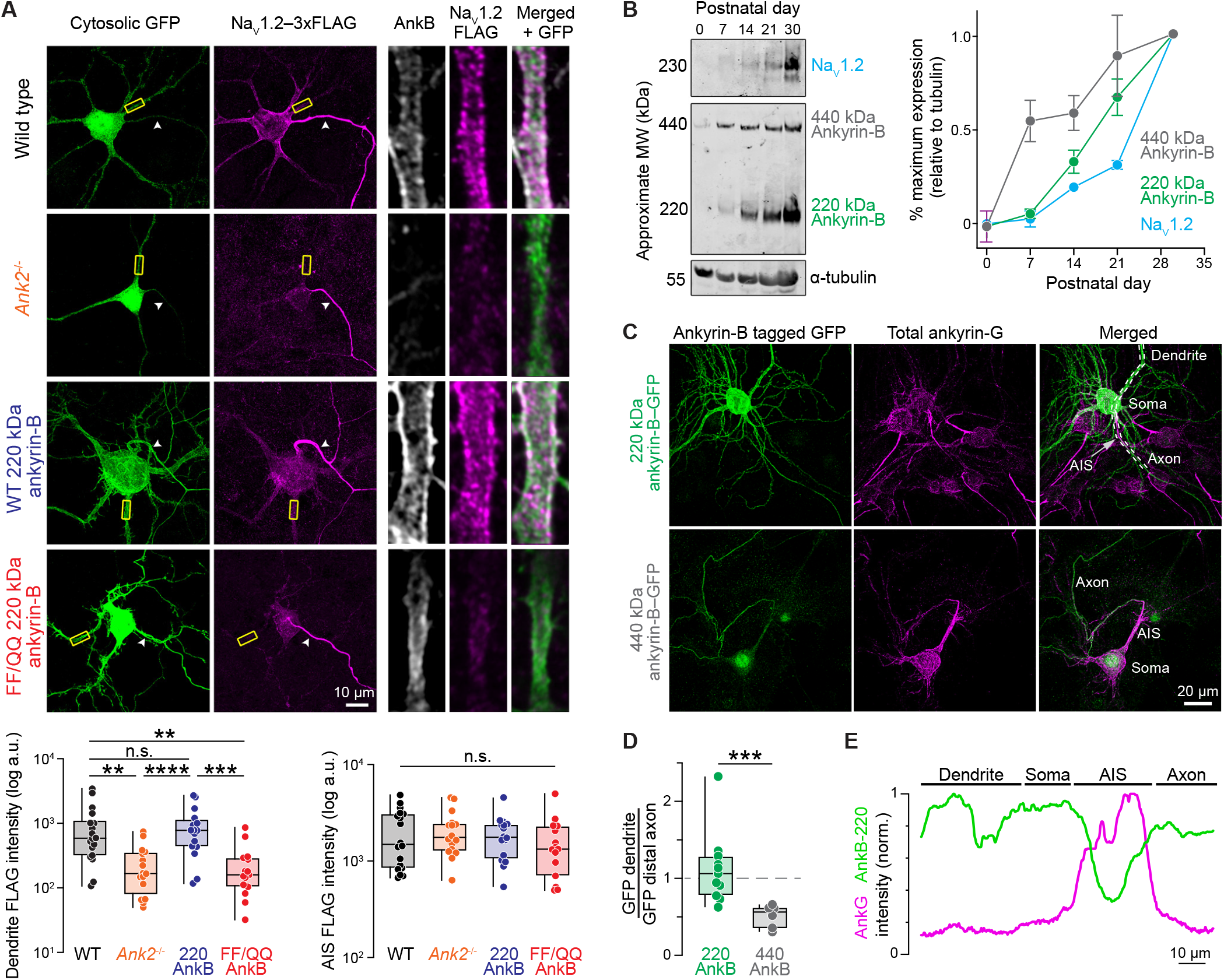
220 kDa ankyrin-B scaffolds Na_V_1.2 to the dendritic membrane. **A**. Top left: Confocal images of DIV21 WT or total *Ank2-*null cultured neocortical neurons co-transfected with Na_V_1.2-3xFLAG-IRES-eGFP or Cre-2A-BFP and rescued with WT or FF/QQ mutant 220 kDa ankyrin-B. Cells were immunostained with antibodies against ankyrin-B (white), FLAG (magenta), and GFP (green). Top right: zoomed images of dendrite labeled with yellow box. Bottom left: Quantification of mean fluorescence intensity of Na_V_1.2-3xFLAG in dendrites. Circles represent individual neurons. (WT: 0.44 ± 0.06, n = 18 cells; *Ank2*^*-/-*^: 0.12 ± 0.02, n = 16 cells; *Ank2*^*-/-*^ + WT 220 kDa AnkB: 0.54 ± 0.1, n = 16 cells; *Ank2*^*-/-*^ + FF/QQ 220 kDa AnkB: 0.15 ± 0.02, n = 15 cells). WT vs. *Ank2*^*-/-*^ **p = 0.0012, *Ank2*^*-/-*^ vs 220 AnkB ****p < 0.0001, WT vs. FF/QQ AnkB **p = 0.0034, 220 AnkB vs. FF/QQ AnkB ***p = 0.0002. Holm-Šídák multiple comparisons test. Bottom right: Quantification of mean fluorescence intensity of Na_V_1.2-3xFLAG in the AIS. No significant differences. Holm-Šídák multiple comparisons test. **B**. Left: Western blot of WT mouse neocortical lysates at P0, P7, P14, P21, and P30. Blots were probed with antibodies to endogenous total ankyrin-B (labeling both 440 kDa (grey) and 220 kDa (green) ankyrin-B isoforms), Na_V_1.2 (cyan), and -tubulin. Right: Quantification of western blot normalized to maximal expression percentage at P30. N = 3 mice per age. Data shown as mean SEM. **C**. Confocal images of WT cultured neurons transfected with 220 kDa AnkB-GFP or 440 kDa AnkB-GFP and fixed at DIV21. Neurons were immunostained with antibodies against endogenous ankyrin-G (magenta) and GFP (green). **D**. Quantification of mean fluorescence intensity (a.u.) ratio of dendritic GFP to distal axon GFP of 220 kDa ankyrin-B (green) versus 440 kDa ankyrin-B (grey) in (C). (220 AnkB: 1.12 ± 0.14, n = 11 cells; 440 AnkB: 0.51 ± 0.06, n = 6 cells). ***p = 0.0003. Mann-Whitney test. **E**. Plot profiles of mean fluorescence intensity (a.u.) of transfected 220 kDa ankyrin-B-GFP versus endogenous total ankyrin-G across different neuronal domains in (C). White dotted line indicates regions measured.

Ankyrin-B is highly homologous with ankyrin-G, especially throughout the ankyrin repeat domain, which contains the canonical Na_V_ binding site (Cai and Zhang, 2006). Therefore, we examined whether ankyrin-B requires this same sequence to localize Na_V_1.2. We generated a double mutant (F131Q, F164Q) in the ankyrin repeats of ankyrin-B, which has been shown previously to reduce binding affinity between ankyrins and Na_V_1.2 by more than 40-fold (Wang et al., 2014). Knockout-and-rescue with the FF/QQ mutant ankyrin-B failed to scaffold Na_V_1.2 to the dendrites, demonstrating the importance of this site for the proper localization of Na_V_1.2 to adult neocortical dendrites (**Figure 2A**). Importantly, Na_V_1.2 clustered appropriately to the AIS in *Ank2* null and FF/QQ ankyrin-B mutant neurons to similar levels as endogenous WT and rescue ankyrin-B conditions (**Figure 2A**). This suggests that the presence of Na_V_1.2 in the dendrites is not an artifact of Na_V_1.2-3xFLAG overexpression and requires ankyrin-B. In addition, the localization patterns of Na_V_1.2-3xFLAG across these conditions are consistent with previous studies using two distinct methods to label endogenous Na_V_1.2 (Gao et al., 2019; Liu et al., 2021a).

Ankyrin-B is expressed as two splice variants in the brain that both contain the conserved Na_V_-binding site in the N-terminal ankyrin repeats (Wu et al., 2015). They consist of a 220 kDa isoform and a giant 440 kDa isoform, which arises from alternative splicing of a single large exon in the middle of the gene (Cunha et al., 2008). The 440 kDa ankyrin-B has been shown to be the primary isoform in the distal axon (Creighton et al., 2021; Lorenzo et al., 2014; Yang et al., 2019), but which splice variant functions in the dendrites is less clear. Western blot analysis of mouse neocortical lysates ranging from P0 to P30 revealed parallel increases in the expression levels of Na_V_1.2 and ankyrin-B throughout development (**Figure 2B**). The marked increase in Na_V_1.2 expression in mouse neocortex from P7 to P30, when the dendritic arbors are elaborating, further highlights the importance of Na_V_1.2 in adult neuronal excitability outside of its early-stage role in the AIS. Expression of the 220 kDa ankyrin-B, which has been shown to be more prominent later in development (Lorenzo et al., 2014; Smith and Penzes, 2018), correlated more closely with Na_V_1.2 expression than the 440 ankyrin-B, suggesting that the 220 kDa ankyrin-B may be the main isoform that functions in the dendrites (**Figure 2B**). To evaluate the localization patterns of the 220 versus the 440 kDa ankyrin-B, we cloned full-length 220 kDa ankyrin-B-GFP and 440 kDa ankyrin-B-GFP. We expressed each construct in cultured neocortical neurons, and immunostained with antibodies against GFP and ankyrin-G. As previously shown, the 440 kDa ankyrin localized almost exclusively to the distal axon (**Figure 2C-D**). By contrast, the 220 kDa isoform localized throughout the entire dendritic arbor as well as the soma and the axon (**Figure 2C-E**). Quantification of mean fluorescence intensity showed the 220 kDa ankyrin-B is distributed across dendrites and the distal axon, whereas the 440 kDa ankyrin-B is primarily in the distal axon (**Figure 2C-E**). Overall, these data indicate that the 220 kDa ankyrin-B is the predominant ankyrin that targets Na_V_1.2 channels to pyramidal cell dendrites.

### Ankyrin-B directly interacts with Na_V_1.2 in adult mouse brain

We next evaluated the molecular basis underlying the interaction between ankyrin-B and dendritic Na_V_1.2. Due to their large size, it is difficult to study direct protein-protein interactions between full-length ankyrins and ion channels (Garrido et al., 2003; Lemaillet et al., 2003). Therefore, we evaluated binding between ankyrin-B-GFP and an epitope-tagged fragment of Na_V_1.2 that contains a highly conserved core nine amino acid motif necessary for ankyrin binding (Na_V_1.2 II-III loop-HA) (Lemaillet et al., 2003) (**Figure 3A**). We co-transfected these constructs into HEK293 cells and performed a proximity ligation assay (PLA), which reports whether ankyrin-B and the Na_V_1.2 II-III loop are associated within 10 nanometers of one another. Using PLA, we found that full-length ankyrin-B readily complexes with the Na_V_1.2 II-III loop in HEK293 cells (**Figure 3B**). To assess direct binding between ankyrin-B and Na_V_1.2, we then transfected HEK293 cells with a mutant Na_V_1.2 II-III loop (termed Δ9-mutant Na_V_1.2) from which we excised the core 9 amino acid ankyrin-binding motif. Expression of the Δ9-mutant Na_V_1.2 completely abolished PLA signal, indicating the importance of this motif for binding between ankyrin-B and Na_V_1.2 (**Figure 3B**). Of note, deletion of the Δ9 sequence in the Na_V_1.2-3xFLAG construct did not affect channel biophysical properties when compared to WT Na_V_1.2 recordings in HEK cells (**Figure S1C**). We further validated their interaction by successfully immunoprecipitating ankyrin-B-GFP and Na_V_1.2 II-III loop from HEK293 cells (**Figure 3C**). Again, we failed to detect any Δ9-mutant Na_V_1.2 following immunoprecipitation of ankyrin-B (**Figure 3C**).

**Figure 3:**
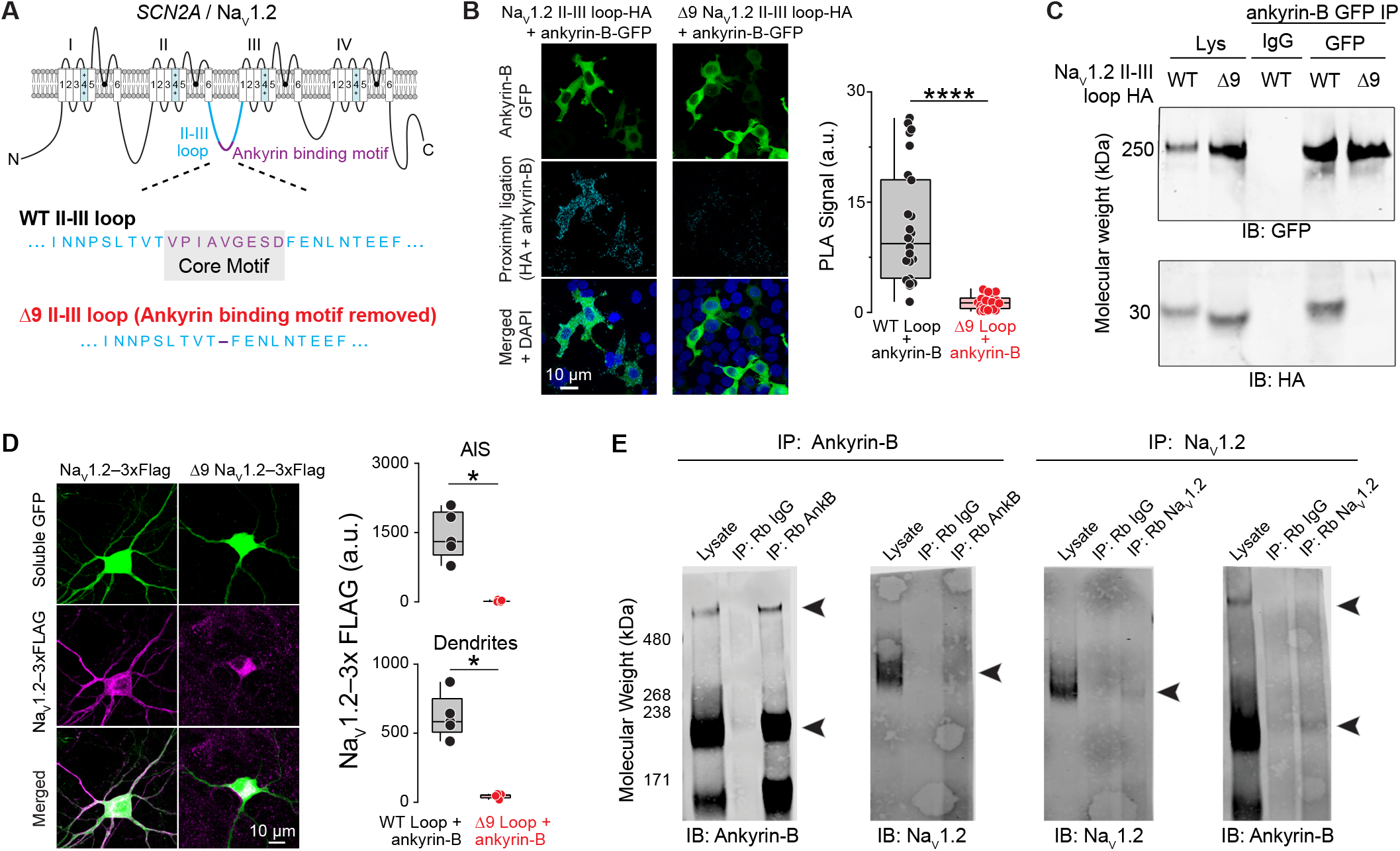
Ankyrin-B directly interacts with Na_V_1.2 in adult mouse brain. **A**. Schematic of Na_V_1.2 highlighting the ankyrin-binding motif (purple) located within the intracellular loop between domains II and III (cyan). Core nine amino acids (D9 motif) within the II-III loop are essential for ankyrin binding. **B**. Left: Representative images of proximity ligation assay (PLA) signal (cyan) between anti-HA and anti-ankyrin-B antibodies from HEK293 cells transfected with ankyrin-B-GFP (green) and HA-tagged Na_V_1.2 II-III loop (left) or the HA-tagged D9 mutant loop (right). Right: Quantification of PLA signal (a.u.) between ankyrin-B and WT Na_V_1.2 II-III loop versus ankyrin-B and D9 Na_V_1.2 II-III loop. (WT: 11.6 ± 1.5, n = 26 cells; D9: 1.4 ± 0.2, n = 24 cells). ****p < 0.0001. Mann-Whitney test. **C**. Co-immunoprecipitation (co-IP) of ankyrin-B-GFP with WT Na_V_1.2 II-III loop or D9 Na_V_1.2 II-III loop. Western blots were probed with antibodies against anti-GFP (to label ankyrin-B-GFP) and anti-HA (to label Na_V_1.2 II-III loop-HA). Non-immune IgG used as a negative control. **D**. Left: Confocal images of cultured neocortical neurons transfected with WT Na_V_1.2-3xFLAG-IRES-eGFP or D9 Na_V_1.2-3xFLAG-IRES-eGFP. Cells were immunostained with anti-GFP (green) and anti-FLAG (magenta) antibodies. Right: Quantification of mean fluorescence intensity (a.u.) of Na_V_1.2-3xFLAG in the AIS (top) and dendrites (bottom). AIS: (WT: 1445 ± 233.5, n = 5 cells; D9: 17.06 ± 15.2, n = 4 cells). *p = 0.016. Mann-Whitney test. Dendrites: (WT: 620 ± 71.0, n = 5 cells; D9: 46.1 ± 9.7, n = 4 cells). *p = 0.016. Mann-Whitney test. **E**. Left: IP of endogenous ankyrin-B and western blot of IP lysates probed with antibodies to ankyrin-B or endogenous Na_V_1.2 from P60-P75 mice. Right: IP of endogenous Na_V_1.2 and western blot of IP lysates probed with antibodies to Na_V_1.2 or ankyrin-B from P60-P75 mice. Black arrows highlight bands of ankyrin-B or Na_V_1.2. Non-immune IgG used as a negative control. Note: 440 kDa ankyrin-B band consistently runs anomalously high as reported previously (Jenkins et al., 2015).

We next tested whether ankyrin-B localized full-length Na_V_1.2 in pyramidal cell dendrites if these channels lacked the ankyrin-binding motif. Cultured neurons were transfected with a plasmid encoding either WT Na_V_1.2-3xFLAG-IRES-GFP or Δ9 Na_V_1.2-3xFLAG-IRES-GFP. At DIV21, WT Na_V_1.2-3xFLAG properly localized to the dendrites as well as the AIS; however, the Δ9-mutant Na_V_1.2-3xFLAG-IRES-GFP, which is unable to interact with ankyrin-B, failed to localize to the dendrites (**Figure 3D**). Since excising the nine amino acid motif prevents all ankyrins from binding, Na_V_1.2-3xFLAG clustering at the AIS was also lost (**Figure 3D**). Additionally, these data with the Δ9-mutant Na_V_1.2 are consistent with those obtained with the FF/QQ mutant ankyrin-B, further confirming that the dendritic localization of Na_V_1.2-3xFLAG is not an artifact of overexpression. The experiments above demonstrate that ankyrin-B is in complex with

Na_V_1.2 in pyramidal neuron dendrites when each protein is overexpressed in cultured neocortical neurons. To determine if this interaction occurs with endogenous ankyrin-B and Na_V_1.2 in native adult neocortex, we immunoprecipitated ankyrin-B using antibodies that detect both the 220 kDa and 440 kDa isoforms. Western blot and immunoblotting with antibodies against Na_V_1.2 revealed ankyrin-B is in complex with Na_V_1.2 in the adult neocortex (**Figure 3E and S2**). Since ankyrin-B is expressed in almost every cell-type within the neocortex (Smith and Penzes, 2018), we wanted to confirm their interaction in pyramidal cell dendrites, where Na_V_1.2 predominantly resides, by immunoprecipitating Na_V_1.2 from adult brain. Immunoprecipitation of Na_V_1.2 resulted in co-IP of the dendritic 220 kDa ankyrin-B (**Figure 2C**), but we did not detect the 440 kDa isoform, which is predominantly axonal (**Figure 3E**). These data provide further evidence that the 220 kDa ankyrin-B is the primary isoform that interacts with Na_V_1.2 in adult mouse brain. Overall, these data demonstrate that endogenous Na_V_1.2 is associated in complex with ankyrin-B in adult neocortex.

### *Ank2* haploinsufficiency impairs dendritic, but not somatic, excitability

Na_V_1.2 channels are expressed on both the somatic and dendritic membrane of mature neocortical pyramidal neurons. Data shown above indicate that ankyrin-B is important for dendritic Na_V_ scaffolding; however, previous reports have demonstrated that both ankyrin-B and ankyrin-G are present on the soma of mature pyramidal neurons (Jenkins et al., 2015; Nelson et al., 2019; Tseng et al., 2015). As such, *Ank2* haploinsufficiency is expected to impair measures of dendritic excitability, but whether somatic excitability is also affected may depend instead on how channels interact with ankyrin-G at the soma. Previously, we used the peak velocity of the rising phase of the AP (peak dV/dt) and AP-evoked dendritic calcium imaging as functional proxies for Na_V_1.2 membrane density in the somatic and dendritic compartments, respectively (Spratt et al., 2019, 2021). While we hypothesized that *Ank2* loss would impair measures of dendritic excitability, we wanted to determine whether it would also affect somatic excitability.

To test how *Ank2* haploinsufficiency affects neuronal AP electrogenesis and propagation, we recorded AP waveform properties from layer 5 (L5) thick-tufted (pyramidal tract) neurons in acute slices of P43-75 *Ank2*^+/ fl^::CaMKIIa-Cre mice, which express Cre in all neocortical pyramidal neurons after ∼P16 (Spratt et al., 2019; Xu et al., 2000) (**Figure 4A-E and S4A**). This allowed for the study of *Ank2* haploinsufficiency in mature cells, a period with a previously defined ASD-associated role for *Scn2a* in pyramidal cell dendrites (Spratt et al., 2019). Western blots of neocortical lysates generated from adult *Ank2*^+/fl^::CaMKIIa-Cre and WT littermates confirmed a significant reduction in the 440 kDa and 220 kDa isoforms of ankyrin-B, without any change in Na_V_1.2 or ankyrin-G expression levels (**Figure S3**). While interleaved experiments in *Scn2a*^+/-^ cells revealed expected reductions in peak somatic dV/dt, *Ank2*^+/fl^ cells were not different than WT (**Figure 4D-E**). In addition, we observed no difference in the FI curves of *Ank2*^+/fl^ neurons compared to WT and *Scn2a*^+/-^ neurons, or in measures of AIS excitability (threshold and AIS-associated peak dV/dt) (**Figure 4B-E**).

**Figure 4:**
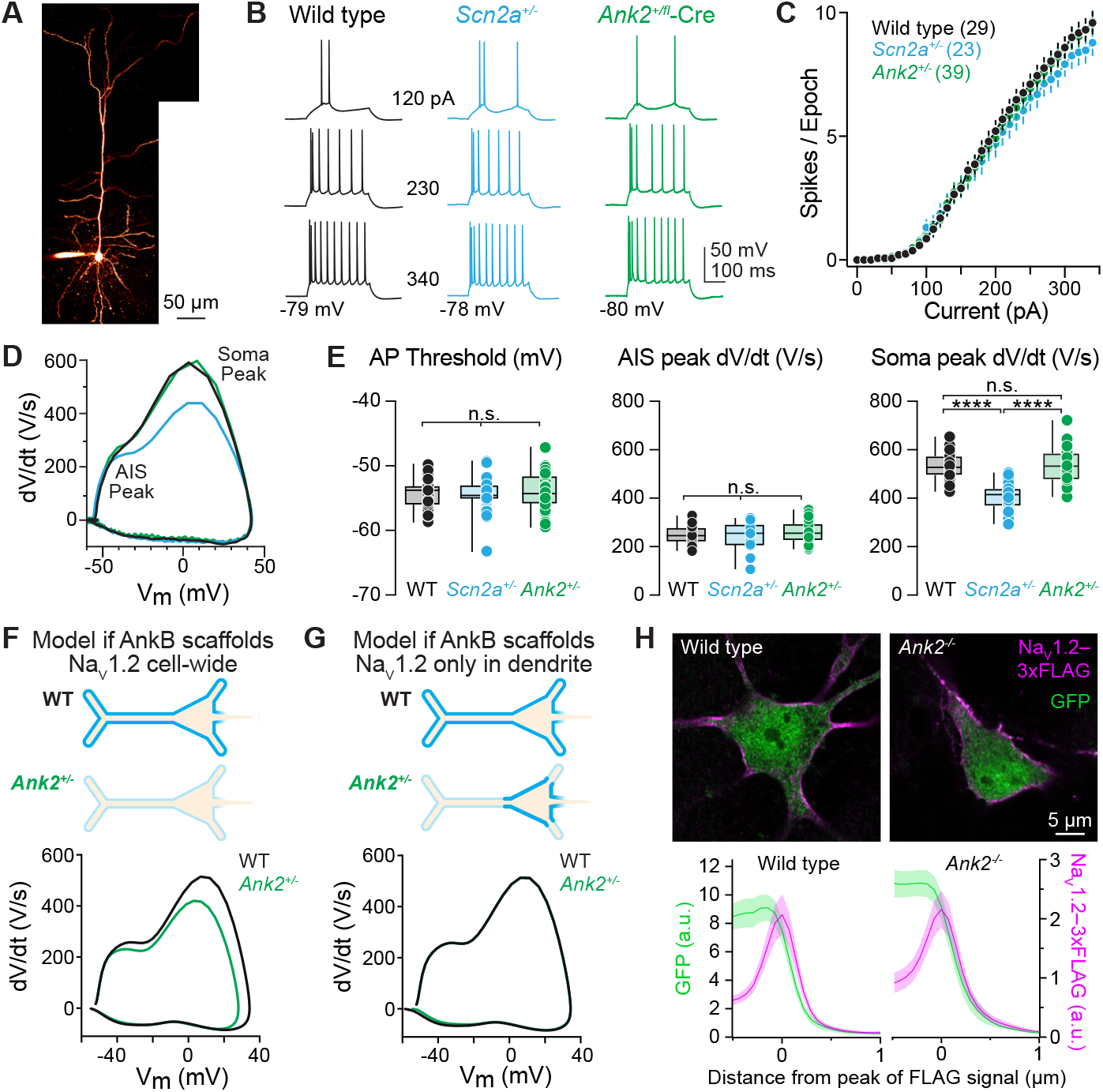
Ank2 haploinsufficiency has no effect on axonal or somatic excitability. **A**. 2-photon laser scanning microscopy (2PLSM) z-stack of a mature layer 5 thick-tufted pyramidal neuron in the mPFC. **B**. APs generated by current injection (0-340 pA, 10 pA intervals, 300 ms) in WT (black), Scn2a^+/-^ (cyan), and Ank2^+/fl^-Cre (green) in P43-75 L5 neurons. **C**. Number of APs (spikes) versus amplitude of current injection as in (B). Firing-rate slope between 0-340 pA: (WT: 0.033 ± 0.002 Hz/pA, n = 29 cells; Scn2a^+/-^: 0.03 ± 0.002 Hz/ pA, n = 23 cells, Ank2^+/fl^-Cre: 0.033 ± 0.001, n = 39 cells). No significant differences. Holm-Šídák multiple comparisons test. **D**. Representative phase-plane plots (dV/dt vs. voltage) of somatic APs from WT (black), Scn2a^+/-^ (cyan), and Ank2^+/fl^-Cre (green) neurons. Different phases of the AP correspond to AP initiation in the AIS (first peak) and soma (second peak). **E**. Left: Quantification of AP threshold (mV) from the first AP evoked by near-rheobase current in P43-75 WT(black), Scn2a^+/-^ (cyan), and Ank2^+/fl^-Cre (green) L5 neurons. Circles represent single cells. (WT: -54.4 ± 0.4 mV, n = 27, Scn2a^+/-^: -54.5 ± 0.4 mV, n = 23 cells, Ank2^+/fl^-Cre: -53.9 ± 0.5 mV, n = 39 cells). No significant differences. Holm-Šídák multiple comparisons test. Middle: AIS AP peak dV/dt (V/s). (WT: 248.7 ± 7.1 V/s, n = 27 cells; Scn2a^+/-^: 242.2 ± 12.3 V/s, n = 23 cells, Ank2^+/fl^-Cre: 257.6 ± 6.8 V/s, n = 39 cells). No significant difference. Holm-Šídák multiple comparisons test. Right: Somatic AP peak dV/dt (V/s). (WT: 537.8 ± 11.3 V/s, n = 27 cells; Scn2a^+/-^: 406.4 ± 11.3 V/s, n = 27 cells, Ank2^+/fl^-Cre: 535.1 ± 10.8 V/s, n = 40 cells). WT vs. Scn2a^+/-^ ****p < 0.0001, Scn2a^+/-^ vs. Ank2^+/fl^-Cre ****p < 0.0001, WT vs. Ank2^+/fl^-Cre p = 0.86 Holm-Šídák multiple comparisons test. **F**. Compartmental model of Na_V_1.2 loss from both somatic and dendritic domains on AP waveform. Top: Schematic of Na_V_1.2 distribution across soma and dendrites in WT neurons compared to loss of Na_V_1.2 in all compartments in Ank2^+/fl^-Cre neurons. Bottom: Phase-plane plots from computational model of AP somatic peak dV/dt changes with WT and heterozygous Na_V_1.2 membrane density. No other channels or parameters were altered except for Na_V_1.2. **G**. Compartmental model of Na_V_1.2 loss from dendritic compartments only. Top: Na_V_1.2 localization at soma and dendrites in WT cells versus loss of Na_V_1.2 in dendrites only in Ank2^+/fl^-Cre neurons as predicted based on empirical recordings of L5 neurons in (A-E). Bottom: Phase-plane plots from computational model showing no change in AP peak somatic dV/dt with WT and heterozygous Na_V_1.2 channel densities in distal dendrites only. Somatic and proximal dendrite Na_V_1.2 maintained at WT levels in both models. **H**. Top: Confocal images of DIV21 WT and Ank2-null cultured neocortical neurons transfected with Na_V_1.2-3xFLAG-IRES-eGFP and immunostained with antibodies to anti-FLAG (magenta) and anti-GFP (green). Bottom: Quantification of somatic Na_V_1.2-3xFLAG and GFP mean fluorescence intensity (a.u.) in WT (left) and Ank2-null (right) neurons.

These empirical data suggest that loss of *Ank2* does not affect somatic or axonal excitability. This result contrasts with previous results in *Scn2a*^*+/-*^ neurons and compartmental models where *Ank2* haploinsufficiency results in similar reductions in Na_V_1.2 density throughout the somatodendritic domain (**Figure 4F**). Instead, models indicate that peak dV/dt is dependent exclusively on somatic Na_V_ density and is insensitive to changes in dendritic channel density (**Figure 4G**), and therefore suggests that Na_V_1.2 densities are at WT levels in *Ank2*^*+/-*^ cells in the soma. This motivated us to re-examine Na_V_1.2-3xFLAG labeling in cultured *Ank2*^*-/-*^ neurons, focusing on somatic regions. Indeed, Na_V_1.2-3xFLAG was localized to the soma in these cells at levels comparable to those found in WT neurons (**Figure 4H**). Thus, ankyrin-G may be the primary ankyrin that localizes Na_V_1.2 to the soma or it is able to compensate for somatic ankyrin-B loss in both heterozygous or homozygous knockout conditions.

While somatic recordings described above suggest that somatic Na_V_ density is not affected by *Ank2* haploinsufficiency, they cannot inform on changes in dendritic excitability, which modeling suggests would still be impaired in *Ank2*^*+/-*^ conditions (**Figure S4B**). To evaluate the effects of ankyrin-B loss on dendritic Na_V_ channel density and excitability, we first examined dendritic Na_V_ function directly with AP-evoked Na imaging using the sodium indicator ING-2 (500 µM), which is sensitive to relatively large changes in sodium concentration (K_D_ = 20 mM) but has higher signal- to-noise than other sodium indicators (Blömer et al., 2021; Filipis and Canepari, 2021; Lipkin et al., 2021). In WT neurons, trains of APs (40 APs at 100 Hz) evoked detectable sodium transients within the first 125 µm of the apical dendrite (**Figure 5A**). These transients were largest proximal to the soma and fell off with increasing distance from the soma. At 25 µm from the soma, Na^+^ transient amplitudes were comparable in WT and *Ank2*^*+/-*^ neurons. In *Scn2a*^*+/-*^ neurons, Na^+^ influx was reduced by 50%, consistent with haploinsufficiency of Na_V_ density. Farther from the soma, however, transients in *Ank2*^*+/-*^ neurons became significantly smaller and exhibited a 40% reduction in Na^+^ influx compared to the WT average (**Figure 5A-B**). This stark change in Na^+^ influx between 25 and 50 microns from the soma corresponds well to the distribution of endogenous ankyrin-G, which can extend into the first tens of microns of dendrite in cultured neurons (**Figure 2C**). Thus, these data are most consistent with a model where ankyrin-G is capable of scaffolding Na_V_s in the soma and proximal dendrite in *Ank2* haploinsufficient conditions, but that ankyrin-B is solely responsible for Na_V_ scaffolding in more distal dendritic domains.

**Figure 5:**
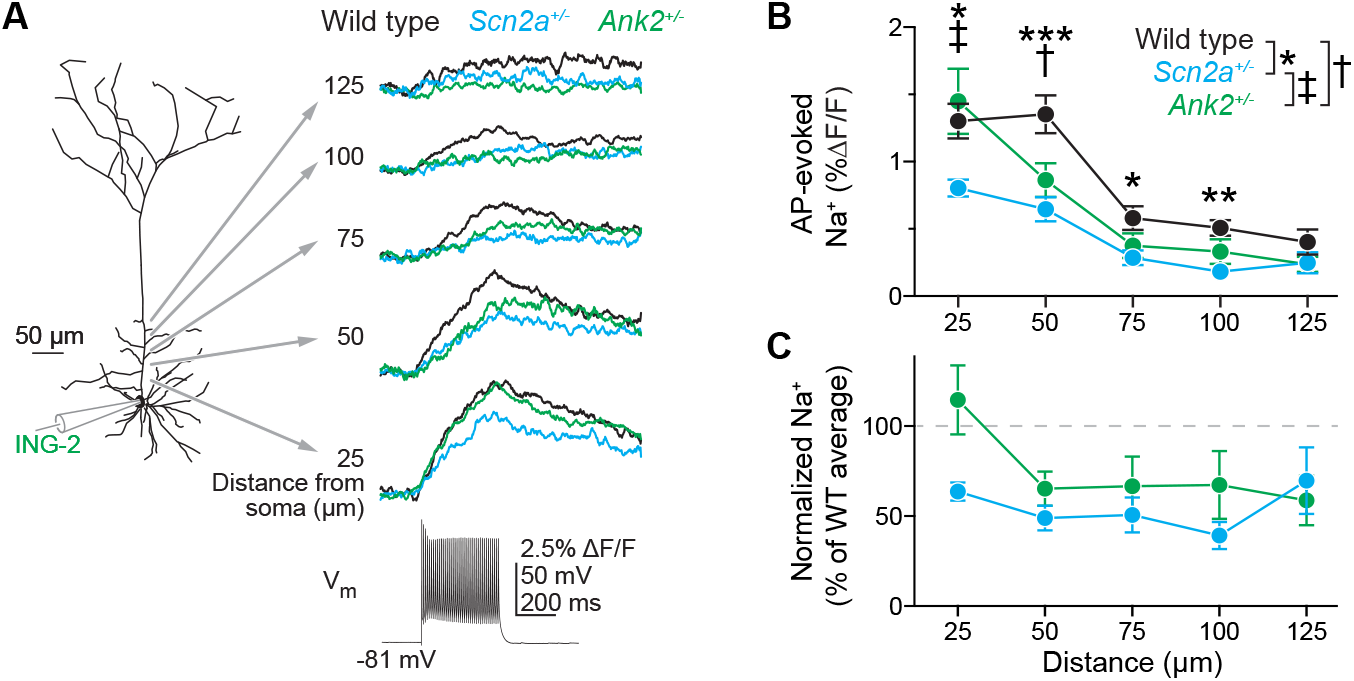
Reduced AP-evoked Na^+^ influx in apical dendrites of Ank2 haploinsufficient neurons. **A**. 2PLSM sodium imaging throughout the apical dendrite of a L5 thick-tufted neuron filled with ING-2 (500 mM) via the recording pipette. Sodium transients were evoked by trains of APs (stimulus: 40x 100 Hz, 2 nA for 2 ms) every 25 microns from the soma in WT (black), *Scn2a*^+/-^ (cyan) and *Ank2*^+/fl^::CaMKIIa-Cre (green) P46-76 mice. Clear signals were detected until 125 microns in WT neurons. **B**. Quantification of AP-evoked Na^+^ transient amplitude versus distance from soma. 25 microns – (WT: 0.013 ± 0.001, n = 11 cells; *Scn2a*^+/-^: 0.008 ± 0.0006, n = 12 cells; *Ank2*^+/ fl^-Cre: 0.015 ± 0.002, n = 8 cells) WT vs. *Scn2a*^+/-^ (*) p = 0.03, WT vs *Ank2*^+/fl^-Cre, p = 0.49, *Scn2a*^+/-^ vs. *Ank2*^+/fl^-Cre (‡) p = 0.01, Holm-Šídák multiple comparisons test. 50 microns – (WT: 0.014 ± 0.001, n = 11 cells; *Scn2a*^+/-^: 0.007 ± 0.001, n = 12 cells; *Ank2*^+/fl^-Cre: 0.009 ± 0.001, n = 8 cells) WT vs. *Scn2a*^+/-^ (***) p = 0.0004, WT vs *Ank2*^+/fl^-Cre (†) p = 0.02, *Scn2a*^+/-^ vs. *Ank2*^+/fl^-Cre p = 0.23, Holm-Šídák multiple comparisons test. 75 microns – (WT: 0.006 ± 0.0009, n = 8 cells; *Scn2a*^+/-^: 0.003 ± 0.0005, n = 9 cells; *Ank2*^+/fl^-Cre: 0.0038 ± 0.0009, n = 8 cells) WT vs. *Scn2a*^+/-^ (*) p = 0.04, WT vs *Ank2*^+/fl^-Cre p = 0.17, *Scn2a*^+/-^ vs. *Ank2*^+/fl^-Cre p = 0.42, Holm-Šídák multiple comparisons test. 100 microns – (WT: 0.005 ± 0.0006, n = 11 cells; *Scn2a*^+/-^: 0.0018 ± 0.0004, n = 11 cells; *Ank2*^+/fl^-Cre: 0.003 ± 0.0009, n = 7 cells) WT vs. *Scn2a*^+/-^ (**) p = 0.001, WT vs *Ank2*^+/ fl^-Cre p = 0.11, *Scn2a*^+/-^ vs. *Ank2*^+/fl^-Cre p = 0.11, Holm-Šídák multiple comparisons test. 125 microns – (WT: 0.004 ± 0.0009, n = 8 cells; *Scn2a*^+/-^: 0.002 ± 0.0008, n = 7 cells; *Ank2*^+/fl^-Cre: 0.002 ± 0.0006, n = 6 cells) WT vs. *Scn2a*^+/-^ p = 0.4, WT vs *Ank2*^+/fl^-Cre p = 0.4, *Scn2a*^+/-^ vs. *Ank2*^+/fl^-Cre p = 0.9, Holm-Šídák multiple comparisons test. Data shown as mean ± SEM. **C**. Na^+^ influx normalized to WT average at each distance and converted to percent change from maximum WT.

Due to the sensitivity of sodium indicators, dendritic Na^+^ influx could be imaged only within 150 microns of the soma. To understand how *Ank2* haploinsufficiency affects excitability in more distal dendritic compartments, we took advantage of the fact that bursts of backpropagating APs (bAPs) reliably engage voltage-dependent calcium channels throughout dendritic arbors of layer 5 prefrontal pyramidal cells (Gulledge and Stuart, 2003; Larkum et al., 1999; Short et al., 2017; Spratt et al., 2019; Stuart and Häusser, 2001). Importantly, this dendritic Ca influx is mediated largely by Ca_V_1 and Ca_V_3 channels, which, in contrast to Ca_V_2.1 and Ca_V_2.2 channels, have not been reported to interact with ankyrin-B (Choi et al., 2019; Kline et al., 2014; McKay et al., 2006; Pérez-Garci et al., 2013). Consistent with previous observations, bursts of APs (a set of 5 spike doublets at 100 Hz) evoked robust calcium transients throughout the apical dendrite of WT neurons, including distal dendritic tuft branches (**Figure 6A**). By contrast, Ca^2+^ transients were markedly reduced in the *Ank2*^+/-^ neurons, mirroring observations made in *Scn2a*^*+/-*^ neurons (Spratt et al., 2019) (**Figure 6A**). Of note, dendritic arborization was unaltered in adult L5 *Ank2*^+/-^ neurons, with no difference in branch number or length compared to WT neurons (**Figure S5**). Taken together, these data demonstrate that *Ank2* and *Scn2a* converge to regulate dendritic, but not somatic, intrinsic excitability.

**Figure 6:**
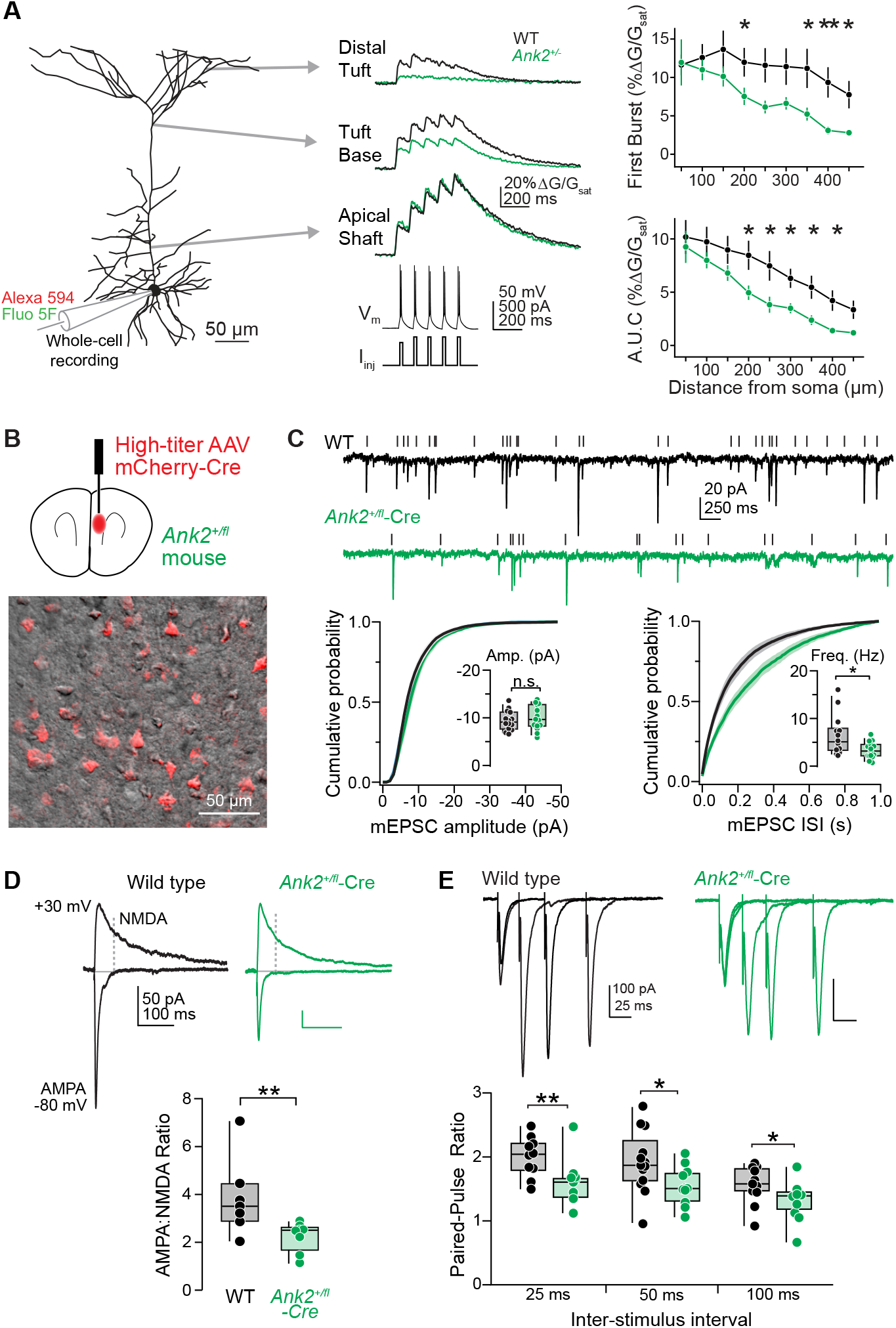
Ank2 haploinsufficiency impairs dendritic backpropagation of APs and excitatory synapse function. **A**. Left: 2PLSM imaging of Ca transients evoked by bursts of AP doublets in L5 thick-tufted pyramidal neuron dendrites in P52-67 WT and Ank2^+/fl^::CaMKIIa-Cre mice. Right: Ca transient amplitude plotted for the first burst (top) and the area under the curve (a.u.c.) of all 5 bursts (bottom) vs. distance. (WT: n=8, Ank2^+/fl^-Cre: n=7). Error bars are mean ± SEM. Mann-Whitney test. *p < 0.05. **B**. Ank2^+/fl^ mice were injected with AAV5-Ef1a-Cre-mCherry virus, transducing pyramidal neurons in the mPFC. 2PLSM single optical sections of mCherry fluorescence (red) overlaid with scanning DIC image (grayscale). **C**. mEPSCs recorded from P54-P60 WT (black) and Ank2^+/fl^- Cre-mCherry-positive (green) neurons. Tick marks denote detected events. Left: Cumulative probability distribution of mEPSC amplitudes. Distributions were generated per cell and then averaged. Insert: average mEPSC amplitude per cell. (WT: -9.4 ± 0.6 pA, n = 14 cells; Ank2^+/fl^-Cre: -10.1 ± 0.7 pA, n = 15 cells) p = 0.5 Mann-Whitney test. Right: Cumulative probability distribution of mEPSC inter-event intervals (IEI). Insert: average mEPSC frequency per cell. (WT: 6.4 ± 1.1 Hz, n = 14 cells; Ank2^+/fl^-Cre: 3.3 ± 0.4 Hz, n = 15 cells) *p = 0.01 Mann-Whitney test. **D**. Top: AMPA-receptor mediated (-80 mV) and mixed AMPA- NMDA (+30) evoked EPSCs from P54-60 WT (black) and Ank2^+/fl^-Cre-mCherry-positive (green) neurons. Dotted line indicates when NMDA component was calculated (50 ms after stimulation onset). Bottom: Quantification of AMPA:NMDA ratio. (WT: 3.9 ± 0.6, n = 7 cells; Ank2^+/fl^-Cre: 2.2 ± 0.2, n = 8 cells) **p = 0.0093. Mann-Whitney test. **E**. Top: Paired-pulse ratio of evoked excitatory inputs at 25, 50, and 100 ms intervals in P50-67 WT (black) and Ank2^+/ fl^::CaMKIIa-Cre (green) mice. Bottom: Summary of PPR grouped by inter-stimulus interval. 25 ms – (WT: 2.0 ± 0.1, n = 10 cells; Ank2^+/fl^-Cre: 1.6 ± 0.1, n = 11 cells) **p = 0.0079. Mann-Whitney test. 50 ms – (WT: 1.9 ± 0.1, n = 13 cells; Ank2^+/fl^-Cre: 1.5 ± 0.09, n = 11 cells) *p = 0.03. Mann-Whitney test. 100 ms – (WT: 1.6 ± 0.1, n = 11 cells; Ank2^+/fl^-Cre: 1.3 ± 0.08, n = 12 cells) *p = 0.0317. Mann-Whitney test.

### *Ank2* heterozygous mice demonstrate impaired excitatory synaptic function

In *Scn2a*^*+/-*^ cells, impaired dendritic excitability weakens postsynaptic aspects of excitatory synaptic transmission by reducing the relative number of functionally mature, AMPA-containing synapses (Spratt et al., 2019). We hypothesized that attenuated bAP in *Ank2*^+/-^ dendrites would result in similar synaptic deficits as observed in *Scn2a*^+/-^ mice. To test this, we evaluated pre- and postsynaptic components of synaptic function in *Ank2*^*+/fl*^ mice injected with a Cre-expression adeno-associated virus (AAV-EF1a-Cre-mCherry, injections at P30, experiments at P52-60) (**Figure 6B**). Whole-cell voltage-clamp recordings of miniature excitatory and inhibitory postsynaptic currents (mEPSC, mIPSC) revealed a 40% reduction in mEPSC frequency with no change in mEPSC amplitude (**Figure 6C**). mIPSC frequency and amplitude were unaffected (**Figure S6**). These data are similar to those obtained in *Scn2a*^*+/-*^ cells and are consistent with a reduction in the number of functionally mature, AMPA-containing excitatory synapses; however, there could also be changes in release probability. To test this possibility, we examined excitatory transmission evoked via local electrical stimulation. We found that the AMPA:NMDA ratio was reduced in *Ank2*^+/-^-Cre neurons, consistent with previous observations in *Scn2a*^+/-^ conditions (**Figure 6D**); however, paired-pulse ratio (PPR), which often is inversely proportional to release probability, was also altered, an effect not observed in *Scn2a*^*+/-*^ conditions. Interestingly, we observed a decrease, rather than an increase, in PPR, suggesting that release probability had increased despite an overall reduction in mEPSC frequency (**Figure 6E**). This suggests that ankyrin-B has differential roles in dendritic and axonal compartments and that impaired dendritic excitability results in an increased proportion of AMPA-lacking synapses and a reduction in mEPSC frequency. This large effect of silent synapses may mask the expected increases in mEPSC frequency due to increased release probability.

To test this, we isolated the postsynaptic contributions of ankyrin-B by injecting a dilute AAV-EF1a-Cre-mCherry virus into the mPFC of *Ank2*^*+/ fl*^ mice at P30 (**Figure 7A**). In these conditions, cell-autonomous effects of *Ank2* haploinsufficiency in dendrites could be assessed by patching one of the few mCherry positive neurons that largely receive input from largely mCherry-negative (e.g., WT) inputs (**Figure 7B**). In these conditions, we found a significant reduction in AMPA:NMDA ratio in *Ank2*^*+/fl*^ neurons with no change in PPR (**Figure 7C-D**). Taken together, these results indicate that, postsynaptically, ankyrin-B is critical for scaffolding of dendritic Na_V_s and that *Ank2* haploinsufficiency phenocopies dendritic deficits observed in *Scn2a*^*+/-*^ neurons. Additional functions for *Ank2* are present in axons, where its loss increases release probability. Thus, in neocortex, *Ank2* haploinsufficiency converges with *Scn2a* haploinsufficiency in pyramidal cell dendrites, but has points of divergence in axons and the soma.

**Figure 7:**
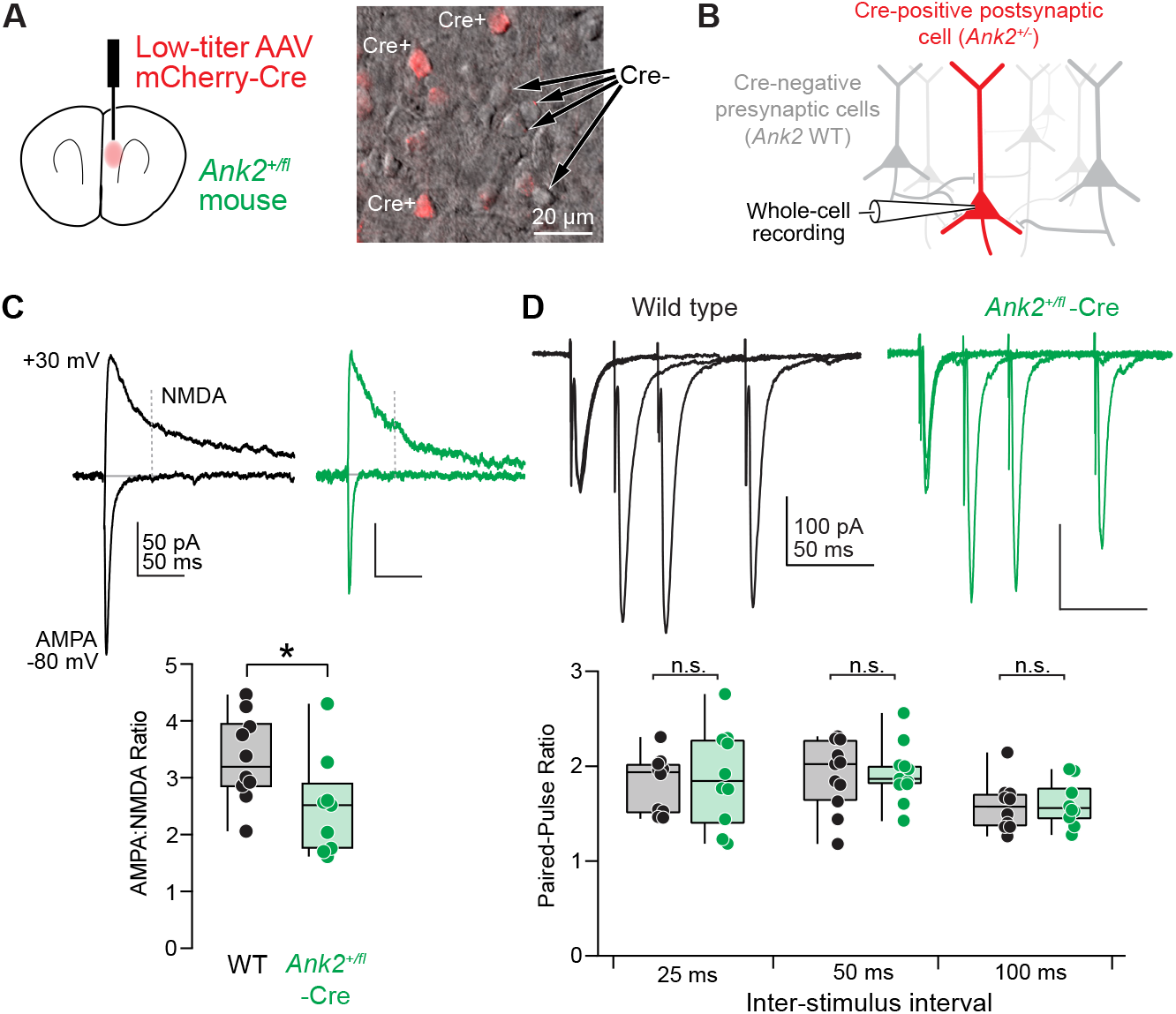
Cell-autonomous *Ank2* haploinsufficiency results in postsynaptic excitatory synaptic function. **A**. Left: Neurons were sparsely transduced by injecting a diluted AAV5-Ef1a-Cre-mCherry virus in mPFC of P30 *Ank2*^*+/fl*^ neurons. Right: 2PLSM single optical sections of mCherry fluorescence (red) overlaid with scanning DIC image (grayscale) showing Ef1a-Cre-mCherry expression in a subset of *Ank2*^*+/fl*^ L5 pyramidal neurons. **B**. Schematic demonstrating *Ank2*^*+/*fl^-Cre-mCherry neuron targeted for whole-cell recording (red). Presynaptic Cre-neurons are WT (gray). **C**. Top: AMPA:NMDA ratio of evoked EPSCs from P52-73 WT (black) and *Ank2*^*+/*fl^-Cre-mCherry (green) neurons. Bottom: Summary of AMPA:NMDA ratio. (WT: 3.3 ± 0.2, n = 10 cells; *Ank2*^+/fl^-Cre: 2.5 ± 0.3, n = 9 cells) *p = 0.02. Mann-Whitney test. **D**. Top: Paired-pulse ratio of evoked excitatory inputs at 25, 50, and 100 ms intervals in P52-73 WT (black) and *Ank2*^*+/*fl^-Cre- mCherry (green) neurons. Bottom: PPR grouped by inter-stimulus interval. 25 ms – (WT: 1.8 ± 0.1, n = 10 cells; *Ank2*^+/ fl^-Cre: 1.9 ± 0.2, n = 10 cells) p > 0.99. Mann-Whitney test. 50 ms – (WT: 1.9 ± 0.1, n = 11 cells; *Ank2*^+/fl^-Cre: 1.9 ± 0.1, n = 11 cells) p = 0.7. Mann-Whitney test. 100 ms – (WT: 1.6 ± 0.9, n = 10 cells; *Ank2*^+/fl^-Cre: 1.6 ± 0.07, n = 10 cells) p = 0.6. Mann-Whitney test.

## DISCUSSION

Ankyrin-B variants have long been known to contribute to cardiac dysfunction through their function in scaffolding a range of membrane pumps, ion exchangers, and receptors (Mohler et al., 2003). Here, we provide evidence that ankyrin-B (*ANK2*) functions as the primary scaffold for Na_V_1.2 (*SCN2A*) in the dendrites of neocortical pyramidal neurons. Haploinsufficiency of *Ank2* in prefrontal neocortical neurons caused dendritic excitability and synaptic deficits, due to reduced Na_V_ channel density within the dendritic membrane, which phenocopies *Scn2a* haploinsufficient conditions. These findings establish a direct, convergent mechanism between two major ASD-associated genes and add to a growing body of literature demonstrating dysfunction in dendritic excitability in ASD (Brager and Johnston, 2014; Brandalise et al., 2022; Johnston et al., 2016; Nelson and Bender, 2021).

### Subcellular patterning of ankyrins

While ion channels and ankyrin scaffold interactions have been studied extensively in axonal domains (Bender and Trussell, 2012; Huang and Rasband, 2018; Kole and Stuart, 2012; Leterrier, 2018; Nelson and Jenkins, 2017), the mechanisms governing dendritic localization of Na_V_s have received far less attention. Data here support a model where ankyrin-G and ankyrin-B have largely distinct roles in Na_V_ scaffolding, with ankyrin-G localized to excitable parts of the axon (e.g., AIS and nodes of Ranvier), and ankyrin-B localized to other regions, including dendrites. We show that ankyrin-B is critical for scaffolding Na_V_1.2 in this domain, and does so along the dendritic shaft membrane. Consistent with this, complete knockout of ankyrin-B in cultured neurons eliminates dendritic Na_V_1.2 immunostaining (**Figure 1**), and *Ank2*^*+/-*^ conditions decrease AP-evoked dendritic Na influx by ∼50% (**Figure 5**). Interestingly, ankyrin-G is known to localize within spines, but does so in support of synapse function rather than Na_V_ scaffolding (Smith et al., 2014). Thus, these data are consistent with the hypothesis that dendritic sodium channels are scaffolded exclusively by ankyrin-B, and that heterozygous or homozygous loss of ankyrin-B cannot be compensated for by other ankyrins.

Intriguingly, one place where ankyrin-B and ankyrin-G overlap, and appear to serve compensatory Na_V_ scaffolding roles, is at the somatic membrane. Here, we observed a disconnect between *Ank2*^*+/-*^ and *Scn2a*^*+/-*^ conditions, as *Ank2*^*+/-*^ neurons did not show a decrease in peak AP velocity, which is one hallmark of *Scn2a*^*+/-*^ neurons (**Figure 4**) (Spratt et al., 2019, 2021). This indicates that somatic Na_V_s can be scaffolded at the soma at WT levels in *Ank2*^*+/-*^ conditions, either by ankyrin-G, or by preferential recruitment of extant ankyrin-B to the soma over other compartments. In support of the former, we found that Na_V_1.2-3xFLAG constructs were still present at the soma in *Ank2*^*-/-*^ cultured neurons, despite being absent from dendrites (**Figure 4**). Methods to label each ankyrin, their splice variants, and Na_V_ subtypes scaffolded to each of these compartments in neurons will be useful to unravel these complexities.

Taken together, these findings suggest that both ankyrins can scaffold somatic Na_V_s, but why they fail to compensate for one another outside the soma remains unclear. One explanation may lie in how ankyrin-G and ankyrin-B are themselves localized to different neuronal compartments. Previous studies have shown that ankyrin-G requires the post-translational modification *S*-palmitoylation for its membrane association at the AIS (He et al., 2012). *S*-palmitoylation is a process mediated by a family of 23 palmitoyl acyl transferases (zDHHC PATs) that covalently adds a 16-carbon fatty acid chain to a conserved cysteine 70 (C70) that resides within the ankyrin repeats (Globa and Bamji, 2017). Two of these PATs, zDHHC5 and zDHHC8, are known to palmitoylate ankyrin-G at the AIS (He et al., 2012), but whether ankyrin-B is palmitoylated in the dendrites has not been determined. Alternatively, ankyrin localization to discrete domains may be driven by autoinhibition mechanisms from unstructured C-terminal ankyrin domains, which have been shown to play an important role in restricting ankyrin activity throughout the cell (Chen et al., 2017).

In addition to differential localization of ankyrin family members, individual splice variants of ankyrin-G and ankyrin-B have unique localization patterns and functions. Alternative splicing of *ANK2* gives rise to two main isoforms of ankyrin-B in the brain: a canonical 220 kDa isoform and a larger 440 kDa splice variant, which contains a single 6.4-kb neuron-specific exon within the middle of the gene (Chan et al., 1993; Kunimoto, 1995; Kunimoto et al., 1991). Here, we found that the 220 kDa ankyrin-B is expressed throughout dendrites, whereas 440 kDa isoform was dominant in distal axons (**Figure 2**). Using conditional *Ank2* alleles, we removed *Ank2* from prefrontal pyramidal cells to isolate cell-autonomous, postsynaptic roles of the 220 kDa isoform, and found that it is capable of scaffolding Na_V_1.2 through direct interaction. This impaired dendritic excitability and excitatory synaptic function in ways that converged with those observed in *Scn2a*^*+/-*^ neurons (Spratt et al., 2019). While these effects likely contribute to ASD etiology, ASD-associated variants in *ANK2* cannot only affect the 220 kDa ankyrin-B, as the 440 kDa isoform will also be impacted. Most variants in *ANK2* fall in this category, resulting in heterozygous expression and haploinsufficiency of both the 220 and 440 kDa isoforms in humans (Fu et al., 2021; Satterstrom et al., 2020). Nevertheless, three ASD-associated variants have been identified in the neuronal-specific exon in *ANK2* that is only found within the 440 kDa isoform (Yang et al., 2019). Here, we found that a modest decrease in paired-pulse ratio was likely attributed to heterozygous loss of axonal *Ank2*, similar to reports examining a frameshift variant at R2608 that specifically affect the 440 kDa isoform (Yang et al., 2019). How axonal ankyrin-B heterozygosity alters short-term plasticity is unclear, but in the heart, *Ank2*^*+/-*^ enhances calcium release from intracellular stores (Popescu et al., 2016). Such mechanisms, if engaged at axonal boutons, could affect neurotransmitter release (Galante and Marty, 2003). Therefore, changes in the 440 kDa ankyrin-B, which is predominantly found within the distal axon, could lead to altered presynaptic neurotransmitter release.

### Subcellular patterning of Na_V_s

AP waveform analysis in *Scn2a*^*+/-*^ and *Scn2a*^*-/-*^ neurons indicate that neocortical pyramidal cell somata are enriched with equal densities of Na 1.2 and Na 1.6 (Spratt et al., 2019, 2021). Based on these observations, we originally assumed that dendrites would also express similar ratios of Na_V_1.2 and Na_V_1.6. However, several observations made here suggest instead that neocortical dendrites are enriched predominantly with Na_V_1.2. First, both *Scn2a*^*+/-*^ and *Ank2*^*+/-*^ dendrites exhibited roughly 50% reductions in AP-evoked Na^*+*^ transients (**Figure 5**). This is most consistent with a single class of Na_V_s, since the alternative—haploinsufficiency in one of two equal sized pools—should have reduced signals by only 25%. Second, both *Scn2a*^*+/-*^ and *Ank2*^*+/-*^ conditions result in similar impairments in AP-evoked dendritic Ca transients, AMPA:NMDA ratio, and mEPSC frequency (**Figure 6**). This again supports the hypothesis that haploinsufficiency of either gene results in comparable impairments in dendritic Na_V_ electrogenesis. Ultimately, new approaches, including freeze-fracture immuno-electron microscopy or super-resolution imaging of epitope-tagged channels (Gao et al., 2019; Liu et al., 2021b; Lorincz and Nusser, 2010), will be needed to definitively determine Na_V_ distribution in distal dendrites of neocortical pyramidal cells.

### Convergent dendritic dysfunction in ASD

Dysfunction in layer neocortical pyramidal cells has long been implicated in ASD (Willsey et al., 2013). These neurons have a unique structure, with a set of basal and apical dendrites that have arborized to receive distinct inputs from local and long-range sources, respectively (Ramaswamy and Markram, 2015). Dendritic integration, including supralinear dendritic spikes in the apical tuft that occur independent of or coincident with activity in the somatic region, are thought to support several computations, including sensory integration, coincident binding of local and long-range input streams, and top-down modulation of cortical processing (Aru et al., 2020; Branco and Häusser, 2010; Branco et al., 2010; Gidon et al., 2020; Major et al., 2013; Markram et al., 2015; Smith et al., 2013). Indeed, these aspects of dendritic excitability are critical for detection of sensory stimuli and behavioral thresholds (Murayama et al., 2009; Takahashi et al., 2016, 2020; Xu et al., 2012), and are some of the first neuronal signals affected with anesthesia-induced loss of consciousness (Suzuki and Larkum, 2020).

Given these major roles in higher-order cortical processing, it is likely that dysfunction in dendritic integration is an important contributor to neurodevelopmental disorders like ASD and intellectual disability (ID). Consistent with this, *SCN2A* haploinsufficiency, which principally interferes with dendritic excitability, unambiguously confers substantial risk for ASD (Ben-Shalom et al., 2017; Fu et al., 2021; Sanders et al., 2012; Satterstrom et al., 2020). Here, we show that *ANK2* haploinsufficiency converges with *SCN2A* to affect the same dendritic excitability mechanisms. This convergence is direct as ankyrin-B is the obligate scaffold for dendritic Na 1.2, resulting in similar deficits in AP-evoked dendritic Na^*+*^ and Ca^2*+*^ influx, and similar deficits in excitatory synaptic function. Emerging evidence indicates that other ASD-associated genes may similarly affect neocortical pyramidal cell dendritic integration, either via regulation of Na_V_s (Brandalise et al., 2022), other channels that influence dendritic excitability (Bock and Stuart, 2016; Gonzalez et al., 2022; Harnett et al., 2013, 2015; Magee and Johnston, 1995; Shah, 2014), or changes in excitation that promote dendritic non-linear events or inhibition that limits such activity (Gidon and Segev, 2012; Megías et al., 2001; Ujfalussy et al., 2015; Wilson et al., 2012; Zhang et al., 2013). These effects can be overt, with ASD-associated variants directly affecting genes encoding dendritic or synaptic proteins in question, or covert, with ASD-associated variants instead affecting gene regulatory elements that in turn alter protein expression (reviewed in Nelson and Bender, 2021).

Na_V_1.2 is critical for dendritic excitability throughout life. Conditionally-induced heterozygosity of *Scn2a* late in development results in identical impairments in dendritic and synaptic function as observed in constitutive *Scn2a* heterozygotes (Spratt et al., 2019). Excitatory synapses appear similar to those found in immature neurons, with relatively small spine heads and low AMPA:NMDA receptor ratios, suggesting that synapses may be maintained in an immature, pre-critical period state (Spratt et al., 2019). Indeed, restoration of near WT levels of *Scn2a*, either via Cre-induced genetic rescue or CRISPR activator-based upregulation of the residual, functional allele in *Scn2a* heterozygotes, restores dendritic excitability, synapse morphology, and synapse function to WT levels (Tamura et al., 2022). This suggests that restoration of *Ank2* function would have similar benefits, at least in dendritic regions where it actively scaffolds Na_V_1.2. *ANK* genes are too large for traditional gene therapy approaches; however, several other approaches are maturing for gene regulation in neurodevelopmental disorders, with marked progress for a number of genetic conditions (Colasante et al., 2020; Derbis et al., 2021; Han et al., 2020; Tamura et al., 2022; Ure et al., 2016; Weuring et al., 2021; Wolter et al., 2020). In addition, a better understanding of the unique roles of different ankyrin proteoforms in scaffolding and function of various binding partners (Gidon and Segev, 2012; Ujfalussy et al., 2015; Wilson et al., 2012; Zhang et al., 2013) may provide insight into methods that allow ankyrins to better compensate for one another in dendritic compartments. Overall, these data establish a framework that both *Scn2a*^*+/-*^ and *Ank2*^*+/-*^ models are forms of channelopathies contributing to ASD (Ptáíek, 2015), motivating future research on potential convergent impairments in channel and dendritic function associated with other ASD/ID risk genes.

## METHODS

### CONSTRUCTS, ANTIBODIES, AND EXPERIMENTAL MODELS

Human Na_V_1.2-3xFLAG IRES eGFP was generated by Genscript (Piscataway, NJ) by the addition of a short linker (AAARG) and a triple-FLAG epitope (DYKDHDGDYKDHDIDYKDDDDK) to the carboxyl terminus of codon-optimized human Na_V_1.2 IRES eGFP (Ben-Shalom et al., 2017). Δ9 Na_V_1.2-3xFLAG IRES eGFP was created by Genscript by deletion of the necessary nine amino acid core ankyrin-binding motif, described previously (Lemaillet et al., 2003). The HA-tagged Na_V_1.2 II-III loops (wild-type and Δ9) were created by cloning the coding sequence corresponding to amino acids 991-1211 of human Na_V_1.2 into pENTR D-TOPO by polymerase chain reaction. The loops were shuttled into pCSF107mT-GATEWAY-3’-3HA (gift from Todd Stukenberg, Addgene plasmid # 67616) using Gateway LR clonase, according to manufacturer’s directions (Thermo Fisher). Na_V_ β1-V5-2A-DsRed was a generous gift from Dr. Lori Isom (University of Michigan) (Bouza et al., 2021). 220 kDa ankyrin-B-GFP was previously described (Mohler et al., 2002). 440 kDa ankyrin-B-GFP was created by subcloning the additional giant exon from 440 kDa ankyrin-B-Halo (Yang et al., 2019)into 220 kDa ankyrin-B-GFP using BstZ17I and SacII sites. 220 kDa ankyrin-B F131Q/F164Q (Wang et al., 2014) was created by Genscript by site-directed mutagenesis. TagBFP and Cre-2A-TagBFP were previously described (Tseng et al., 2015). All plasmids were sequenced across the entire coding sequence by Sanger sequencing prior to use in experiments.

Lab-generated antibodies to ankyrin-B and ankyrin-G were described previously, including rabbit anti-ankyrin-G C-terminus (Kizhatil et al., 2007), goat anti-ankyrin-G C-terminus (He et al., 2014), rabbit anti-270/480 kDa ankyrin-G (Jenkins et al., 2015), rabbit anti-480kDa ankyrin-G (Jenkins et al., 2015), rabbit anti-ankyrin-B C-terminus (Ayalon et al., 2008), sheep anti-ankyrin-B C-terminus. Specificity of all lab-generated antibodies are confirmed using respective null mouse tissue. In addition, antibodies are tested for ankyrin cross-reactivity by both immunocytochemistry and western blotting in HEK293 cells expressing 220 kDa ankyrin-B-GFP or 190 kDa ankyrin-G-GFP. Commercial antibodies used in these studies include rabbit anti-Nav1.2 (Abcam, ab65163), mouse anti-FLAG-M2 (Sigma, F3165), chicken anti-GFP (Abcam, ab13970), mouse anti-HA epitope (clone 6E2, Cell Signaling Technologies), rabbit anti-HA epitope (clone C29F4, Cell Signaling Technologies), and guinea pig anti-MAP2 (Synaptic Systems, 188-004).

C57BL/6J mice were obtained from Jackson Laboratories (stock #000664). The *Ank2* exon 24^flox/flox^ mouse line was a gift from Dr. Peter Mohler (The Ohio State University) (Roberts et al., 2019). *Scn2a*^*+/-*^ mice were provided by Drs. E. Glasscock and M. Montal (Mishra et al., 2017; Planells-Cases et al., 2000).

### CO-IMMUNOPRECIPITATION

Whole brain was dissected from C57Bl/6J adult mice (P60-75). Each brain was homogenized in 2 ml of reaction buffer (0.3 M sucrose, 10 mM Phosphate, 2 mM EDTA; pH 7.4), mixed with phosphatase inhibitor and protease inhibitor. 500 ul of 20 mM DSP (Lomant’s Reagent) was added to the sample and incubated for 2 hours on ice, after which the crosslinking reaction was quenched by adding 1x Tris to a final concentration of 50 mM and incubated on ice for 15 minutes. Samples were lysed by mixing with lysis buffer (30 mM Tris, 150 mM NaCl, 2 mM EDTA, 1% IGEPAL, 0.5% Sodium Deoxycholate; pH 6.8) and sonicating 20 times at 1-second-long pulses, followed by ultracentrifugation at 100k x g for 30 minutes. Solubilized proteins were then subjected to immunoprecipitation using magnetic beads bound to antibodies (Bio-Rad SureBeads Protein A; rabbit ankyrin-B 1:250, rabbit Nav1.2 1:100, rabbit IgG 1:250). Lysate samples were rotated with the bead mixture overnight at 4°C. Beads were collected the next day, washed 3 times with lysis buffer, and mixed 1:1 with 5x PAGE buffer (5% SDS, 25% sucrose, 50 mM Tris; pH 9, 0,5 mM EDTA) and heated to 68°C for 10 minutes.

### WESTERN BLOT

Samples were separated on a 3.5-17% gradient gel in 1x Tris buffer, pH 7.4 (40 mM Tris, 20 mM NaOAc, and 2mM NaEDTA) with 0.2% SDS. Transfer to nitrocellulose membrane was performed overnight at 300 mA at 4°C in 0.5x Tris buffer with 0.01% SDS. Membranes were blocked with 5% Bovine Serum Albumin (BSA) in TBS at room temperature for 1 hour and incubated in primary antibodies (rabbit ankyrin-B 1:1000; rabbit ankyrin-G 1:1000; rabbit Nav1.2 1:500; mouse a-tubulin 1:1000) diluted in 5% BSA in TBS-T overnight at 4°C. Membranes were washed 3x for 10 minutes with TBS-T and incubated for 1 hr at room temperature with LiCor fluorescent secondaries (1:15,000) in 5% BSA in TBS-T. Membranes were then washed 3x for 10 minutes in TBS-T, 3x for 5 minutes in ddH_2_O) before ^2 2^ being imaged on LiCor Odyssey Clx imager.

### PROXIMITY LIGATION ASSAY

HEK293 cells were obtained from the American Type Culture Collection and maintained in a humidified environment at 37 °C with 5% CO_2_. Cells were cultured in DMEM (Invitrogen #11995) with 10% fetal bovine serum, 100 units/ml penicillin, and 100 units/ml streptomycin. 100,000 cells were plated onto glass-bottomed dishes (Cellvis) and were allowed to attach for four hours. Cells were transfected with 100 ng of each plasmid (Na_V_1.2 II-III loop-3xHA and 220 kDa ankyrin-B-GFP) with Lipofectamine 2000, according to the manufacturer’s protocol. After 16 hours, cells were fixed with 4% paraformaldehyde for 15 minutes, permeabilized with 0.1X Triton X-100 for 10 minutes and blocked with 5% bovine serum albumin and 0.2% Tween-20 in PBS for 30 minutes. Cells were incubated with primary antibodies (mouse anti-HA and rabbit anti-ankyrin-B C terminus) in blocking buffer overnight in a humidified chamber. The next day, cells were washed with PBS containing 0.2% Tween-20 (PBS-T) three times for 10 minutes and incubated with anti-mouse minus and anti-rabbit plus PLA probes. Samples were processed for ligation amplification using red fluorescent nucleotides, and mounting, according to the manufacturer’s protocol (Duolink, Sigma-Aldrich).

### NEOCORTICAL CULTURES, TRANSFECTIONS, AND IMMUNOFLUORESCENCE

Primary neocortex was dissected from postnatal day 0 (P0) mice and treated with 0.25% trypsin and 100 µg/ml DNase I in 2 mL HBSS with 10mM HEPES, then triturated gently through a glass pipette with a fire-polished tip. The dissociated neurons were then plated on 35mm MatTek dishes, precoated with poly-D-lysine and laminin, in 0.5mL of Neurobasal-A medium containing 10% FBS, B27 supplement, 2 mM glutamine, and penicillin/streptomycin. On day *in vitro* 1 (DIV1), the neurons were washed with Neurobasal-A medium and fed with growth media (2.5mL of fresh Neurobasal-A medium containing 1% FBS, B27, glutamine, penicillin/ streptomycin, and 2.5 µg/ml AraC. On DIV3, plasmids were introduced into neocortical neurons through lipofectamine 2000-mediated transfection. In one tube, 500 ng of each plasmid was added to 200 μL of Neurobasal-A, and in a second tube, lipofectamine 2000 (3 μL/ 1 μg plasmid) was added to 200 μL of Neurobasal-A. The two tubes were then mixed and incubated at room temperature for 15 min. The neuronal growth media was then removed from the dishes and saved, and transfection media was added to the neurons. Cells were incubated in transfection media for 1 hr at 37°C. The transfection media was aspirated, cells were washed once with warm Neurobasal-A, and growth media was added back to plates. The cells were maintained in culture until 7 DIV or 21 DIV and fixed for immunofluorescence as described below.

Dissociated neocortical neurons were fixed for 15 minutes at room temperature with 4% paraformaldehyde, followed by permeabilization with 0.2% Triton in 1X PBS pH7.4 for 10 minutes at room temperature. They were then blocked with blocking buffer (5% BSA, 0.2% Tween 20 in 1X PBS pH7.4) at room temperature for 30 minutes. Primary antibodies were diluted in blocking buffer and incubated overnight at 4 °C. The next day, cells were washed at room temperature three times for 15 minutes with PBS containing 0.2% Tween 20. Then the cells were incubated with secondary antibodies diluted in blocking buffer for one hour at room temperature. The cells were washed at room temperature three times for 15 minutes with PBS containing 0.2% Tween 20 and then mounted with ProLong Gold antifade reagent before imaging with confocal microscopy as described below.

### CONFOCAL MICROSCOPY

Samples were imaged on a Zeiss LSM 880 with Airyscan using a 63× 1.4 Plan-Apochromat objective and excitation was accomplished using 405-, 488-, 561-, and 633-nm lasers. Each experiment was repeated at least three independent times. Measurements were taken using Fiji software (Schindelin et al., 2012). Laser power and imaging parameters were kept constant for each immunocytochemistry condition.

### *IN VITRO* CELL ELECTROPHYSIOLOGY

Voltage-clamp recordings were performed at room temperature in standard whole-cell configuration, using Axopatch 700B amplifier and pClamp (version 10, Axon Instruments, FosterCity, CA) and a Digidata 1440A digitizer (Molecular Devices). Sodium current was recorded in the presence of external recording solution containing in mM: 120 NaCl, 4 KCl, 1 MgCl, 1.5 CaCl, 10 HEPES, 45 Glucose and 30 Sucrose (Ph 7.35 with CsOH; osmolality was 300–305 mOsm). For the β1 subunit co-transfection experiments, the external sodium concentration was reduced to 60mM. Fire-polished patch pipettes obtained from borosilicate glass capillary (WPI) which resistance was between 1.5-3.5 MΩ, were filled with an internal solution containing in mM: 10 NaCl, 105 Cs-Aspartate, 10 CsCl, 10 EGTA, 10 HEPES, (pH 7.2 with H_2_SO_2_). To determine sodium current amplitude and voltage dependence of activation, currents were evoked by depolarization for 250 ms to different potentials (from −120 to 30 mV on 5 or 10 mV steps) from holding potential of −80 mV and a hyperpolarizing −120mV, 250 ms pre-pulse. Voltage-dependence of inactivation was determined by applying a 50 ms test pulse of 0 mV after the 250 ms pulses used for voltage dependence of activation. Series resistance was compensated no more than 40%–65% when needed, and leak subtraction was performed by application of a standard P/4 protocol. Signals were low-pass filtered at 10 kHz, and data were sampled at 40 kHz online. Current densities were determined by dividing current amplitude by the cell capacitance (Cm) measured by pClamp software. Normalized conductance and inactivation curves were generated as previously described (Patino et al., 2009).

### COMPARTMENTAL MODELING

A pyramidal cell compartmental model was implemented in the NEURON environment (v7.7) based on the Blue Brain Project thick-tufted layer 5b pyramidal cell (TTPC1) model used in our previous study (Ben-Shalom et al., 2017; Markram et al., 2015; Ramaswamy and Markram, 2015; Spratt et al., 2021). The TTPC1 model was adjusted to include an AIS, and the original Na channels in the TTPC1 model were replaced with Na_V_1.2 and Na_V_1.6 channels in compartments with densities as previously shown (Spratt et al., 2021). For phase plane comparisons, the first AP was evoked with a stimulus of 500 pA intensity (25 ms duration) in each model configuration. Threshold was defined as the membrane potential when dV/ dt exceeds 15 V/s. For AP backpropagation, a single AP was evoked with a 1.2 nA, 8 ms step current applied to the somatic membrane. In model conditions with only Na_V_1.2 contributing Na conductance in the distal apical dendrite, Na_V_1.6 was replaced with Na_V_1.2 after ∼30 microns from soma and total Na_V_1.2 conductance was increased by a factor of 1.9 to match total conductance levels in the mixed Na_V_1.2/Na_V_1.6 model (since Na_V_1.6 voltage dependence is more hyperpolarized). Voltage was recorded from the soma, shaft of the apical dendrite (460 µm from soma), and branch of the apical tuft (975 µm from soma). Conductance densities for sodium channels in different compartments across models were as in Table 1:

**Table 1:** Conductance values for Na_V_1.2 and Na_V_1.6 in WT conditions in all compartments

| Compartment | Conductance (mho/cm<sup>2</sup>) | Model with both Na<sub>v</sub>1.2 and Na<sub>v</sub>1.6 in distal dendrite | Model with Na<sub>v</sub>1.2 alone in distal dendrite |
| --- | --- | --- | --- |
| Soma | Na<sub>v</sub>1.2<br>Na<sub>v</sub>1.6 | 0.04840837<br>0.04840837 | 0.04840837<br>0.04840837 |
| Proximal AIS (peak) | Na<sub>v</sub>1.2<br>Na<sub>v</sub>1.6 | 20<br>28 | 20<br>28 |
| Distal AIS (peak) | Na<sub>v</sub>1.2<br>Na<sub>v</sub>1.6 | 0<br>28 | 0<br>28 |
| Axon | Na<sub>v</sub>1.2<br>Na<sub>v</sub>1.6 | 0<br>0.0983955 | 0<br>0.0983955 |
| Axon nodes | Na<sub>v</sub>1.2<br>Na<sub>v</sub>1.6 | 0<br>2 | 0<br>2 |
| Proximal dendrite | Na<sub>v</sub>1.2<br>Na<sub>v</sub>1.6 | 2.2877E-07<br>2.2877E-07 | 2.2877E-07<br>2.2877E-07 |
| Distal dendrite | Na<sub>v</sub>1.2<br>Na<sub>v</sub>1.6 | 2.2877E-07<br>2.2877E-07 | 8.6932E-07<br>0 |

### *EX VIVO* ELECTROPHYSIOLOGY

All experiments were performed in accordance with guidelines set by the University of California Animal Care and Use Committee. Mice aged P55-75 were anesthetized under isoflurane. Brains were dissected and placed in 4 °C cutting solution consisting of (in mM) 87 NaCl, 25 NaHCO_2_, 25 glucose, 75 sucrose, 2.5 KCl, 1.25 NaH_2_PO_4_, 0.5 CaCl_2_, and 7 MgCl_2_ and bubbled with 5% CO_2_/95% O_2_. Coronal slices 250 µm-thick were obtained that included the medial prefrontal cortex. Slices were then incubated in a holding chamber with sucrose solution for 30 mins at 33 °C, then placed at room temperature until recording. Recording solution consisted of (in mM): 125 NaCl, 2.5 KCl, 2 CaCl_2_, 1 MgCl_2_, 25 NaHCO_3_, 1.25 NaH_2_PO_4_, 25 glucose, bubbled with 5% CO_2_/95% O_2_. Osmolarity of the recording solution was adjusted to approximately 310 mOsm. All recordings were performed at 32-34 °C.

Neurons were identified using differential interference contrast (DIC) optics for conventional visually-guided whole-cell recording, or with two-photon-guided imaging of AAV-EF1a-Cre-mCherry fluorescence overlaid on a scanning DIC image of the slice. Patch electrodes were pulled from Schott 8250 glass (3-4 MΩ tip resistance). For current-clamp recordings, patch electrodes were filled with a K-gluconate-based internal solution that contained (in mM): 113 K-Gluconate, 9 HEPES, 4.5 MgCl_2_, 0.1 EGTA, 14 Tris_2_-phosphocreatine, 4 Na_2_-ATP, 0.3 Tris-GTP; 290 mOSM, pH: 7.2-7.25. For Ca^2+^ imaging, EGTA was replaced with 250 µM Fluo-5F and 20 µM Alexa 594. For voltage-clamp recordings, a CsCl-based internal solution was used that contained (in mM): 110 CsMeSO_2_, 40 HEPES, 1 KCl, 4 NaCl, 4 Mg-ATP, 10 Na-phosphocreatine, 0.4 Na_2_-GTP, 5 QX-314, and 0.1 EGTA; ∼290 mOsm, pH 7.22.

Electrophysiological recordings were collected with a Multiclamp 700B amplifier (Molecular Devices) and a custom data acquisition program in Igor Pro software (Wavemetrics). Current-clamp recordings of action potential waveform were acquired at 50 kHz and filtered at 20 kHz. Pipette capacitance was compensated by 50% of the fast capacitance measure under gigaohm seal conditions in voltage-clamp prior to establishing a whole-cell configuration, and the bridge was balanced. Voltage-clamp experiments were acquired at 10-20 kHz and filtered at 3-10 kHz. Pipette capacitance was completely compensated, and series resistance was compensated 50%. All data were corrected for measured junction potentials of 12 and 11 mV in K-gluconate and Cs-based internals, respectively. Data inclusion was based on previously established metrics (Bender and Trussell, 2009; Bender et al., 2010; Spratt et al., 2019), and includes measures for recording stability and cell health [e.g., stable series resistance of <18 MΩ, stable membrane potential (V_m_), and input resistance (R_in_), with less than 15% change over data collection epochs]. All recordings were made using a quartz electrode holder (Sutter Instrument) to minimize electrode drift within the slice.

All acute slice recordings were made from layer 5b thick-tufted pyramidal tract (PT) neurons in the medial prefrontal cortex. In current-clamp, layer 5b neurons were characterized as those that exhibited a voltage rebound more depolarizing that V_rest_ in response to a strong hyperpolarizing current (-400 pA, 120 ms) that peaked within 90 ms of current offset and depolarizing (300 ms, 20-300 pA) square current pulses from a holding potential of -80 mV (Clarkson et al., 2017). AP threshold, AIS dV/dt and peak dV/dt measurements were determined from the first AP evoked by a step current (300 ms duration; 200-300 pA) delivered to the somatic pipette within the first 2 minutes of establishing the whole-cell recording configuration. AP threshold was defined as the V_m_ when dV/dt first exceeded 15 V/s. AIS peak dV/dt was defined at the saddle point between two positive inflection points in the second voltage derivative that occur during the depolarizing phase of the AP.

Miniature excitatory and inhibitory postsynaptic currents (mEPSCs, mIPSCs) were acquired in voltage-clamp configuration at -80 mV and 0 mV, respectively, in the presence of 10 µM R-CPP and 400 nM TTX. Events were analyzed using a deconvolution-based event detection algorithm within IgorPro (Pernía-Andrade et al., 2012). Detectable events were identified using a noise threshold of 3.5x with a minimum amplitude of 2 pA and a 2 ms inter-event interval. Events were subsequently manually screened to confirm appropriate event detection. Event detection code is available at https://benderlab.ucsf.edu/resources. Cumulative probability distribution of mEPSCs and mIPSCs event intervals were generated per cell and then averaged. Distributions were compared using the Kolmogorov-Smirnov test. A confidence interval of 95% (P < 0.05) was required for values to be considered statistically significant. In experiments measuring AMPA:NMDA ratio and paired-pulse ratio (PPR), EPSCs were evoked using a bipolar glass theta electrode placed in layer 5b ∼200 µm lateral from the recording neuron. AMPA: NMDA ratio was initially measured at -80 mV to assess the AMPA contribution and then at +30 mV to evaluate the NMDA-mediated component in the presence of 25 µM picrotoxin. AMPA was defined as the peak inward current at -80 mV and NMDA as the outward current 50 ms after stimulus onset at +30 mV. PPR was acquired at -80 mV in the presence of 10 µM R-CPP and 25 µM picrotoxin. mEPSCs, mIPSCs, and AMPA:NMDA ratio were collected from *Ank2*^*+/fl*^ mice injected with AAV-EF1a-Cre-mCherry and wild type littermates. Mice were anesthetized with isofluorane and positioned in a stereotaxic apparatus. 500 nL volumes of AAV-EF1a-Cre-mCherry (UNC vector core) were injected into the mPFC of *Ank2*^*+/fl*^ mice (stereotaxic coordinates [mm]: anterior-posterior [AP] +1.7; mediolateral [ML] -0.35; dorsoventral [DV]: -2.6). Experiments were conducted four-week post-injection. Paired-pulse ratio (PPR) was recorded from *Ank2*^+/fl^::CaMKIIa-Cre mice and wild type littermates. For AMPA:NMDA and PPR experiments in spare Cre-expressing animals, AAV-EF1a-Cre-mCherry was diluted 1:3 in saline, then injected into the mPFC of *Ank2*^*+/fl*^ mice.

### TWO-PHOTON IMAGING

Two-photon laser scanning microscopy (2PLSM) was performed as previously described (Bender and Trussell, 2009). A Coherent Ultra II was tuned to 810 nm for calcium imaging, ING-2 sodium imaging, and morphology experiments. Epi- and transfluorescence signals were captured either through a 40x, 0.8 NA objective for calcium imaging or a 60X, 1.0 NA objective for ING-2 imaging paired with a 1.4 NA oil immersion condenser (Olympus). For calcium imaging, fluorescence was split into red and green channels using dichroic mirrors and band-pass filters (575 DCXR, ET525/70 m-2p, ET620/60 m-2p, Chroma). Green fluorescence (Fluo-5F) was captured with 10770-40 photomultiplier tubes selected for high quantum efficiency and low dark counts (PMTs, Hamamatsu). Red fluorescence (Alexa 594) was captured with R9110 PMTs. Data were collected in linescan mode (2–2.4 Δ(G/R)/(G/R)_max_*100, where (G/R)_max_ was the maximal fluorescence in saturating Ca^2+^ (2 mM) (Yasuda et al., 2004). AP backpropagation experiments were performed in 25 M picrotoxin, 10 µM NBQX and 10µM R-CPP. For ING-2 sodium imaging, the epifluorescence filters were removed and the transfluorescence filters were replaced with a single 535/150 bandpass filter (Semrock) and all fluorescence was collected on HA10770-40 PMTs. Maximum intensity image projections are displayed using the ‘‘Red Hot’’ lookup table in FIJI. Full neuronal and dendritic reconstructions were stitched together using pairwise stitching in FIJI before generation of maximum intensity projections.

### QUANTIFICATION AND STATISTICAL ANALYSIS

Statistical analysis was performed and presented using Graphpad Prism 9 software. Data were acquired from both sexes and no sex-dependent differences were found. All data were analyzed blind to genotype. Unless otherwise noted, data are shown as mean min. to max. box and whisker plots. Data were quantified as mean ± standard error in figure legends and statistical tests were noted. Each data point (n) is indicative of individual neurons. Mean values per cell were compared using the Mann-Whitney test for two groups or Holm-Šídák multiple comparisons test was used to compare three or more groups. Results were considered significant at alpha value of p < 0.05 and n.s. indicates not significant. Group sample sizes were chosen based on standards in the field and previous similar experiments conducted by our group.

## Acknowledgements

We are grateful to members of the Jenkins and Bender labs, as well as Drs. L. Isom, M. Roberts, and L Ptáíek for comments and feedback. We further thank Dr. M. Roberts for help with code associated with analyzing immunofluorescence relative to membrane borders. This work was supported by the Rackham Predoctoral Fellowship (JMP), Cellular and Molecular Biology Training Grant NIH T32GM007315 (KJP and CCE), NIH F32MH125536 (ADN), Simon’s Foundation for Autism Research Initiative (SFARI) pilot grant #675594 (PMJ), NIH R01MH126960 (PMJ and KJB), and R01MH125978 (KJB).

## Author Contributions

SJS, KJB, and PMJ conceived the project. ADN, AMC, JMP, LM, RNCF, KJP, CCE, AAS, KDD, HK, KJB, and PMJ designed and performed experiments. ADN, AMC, JMP, LM, RNCF, KJP, CCE, AAS, KDD, HK, KJB, and PMJ analyzed the data. ADN wrote the manuscript. ADN, KJB, and PMJ edited the manuscript.

