## Supplemental Figures for "Physical and functional convergence of the autism risk genes *Scn2a* and *Ank2* in neocortical pyramidal cell dendrites"

Content:

Supplemental Figures 1-6

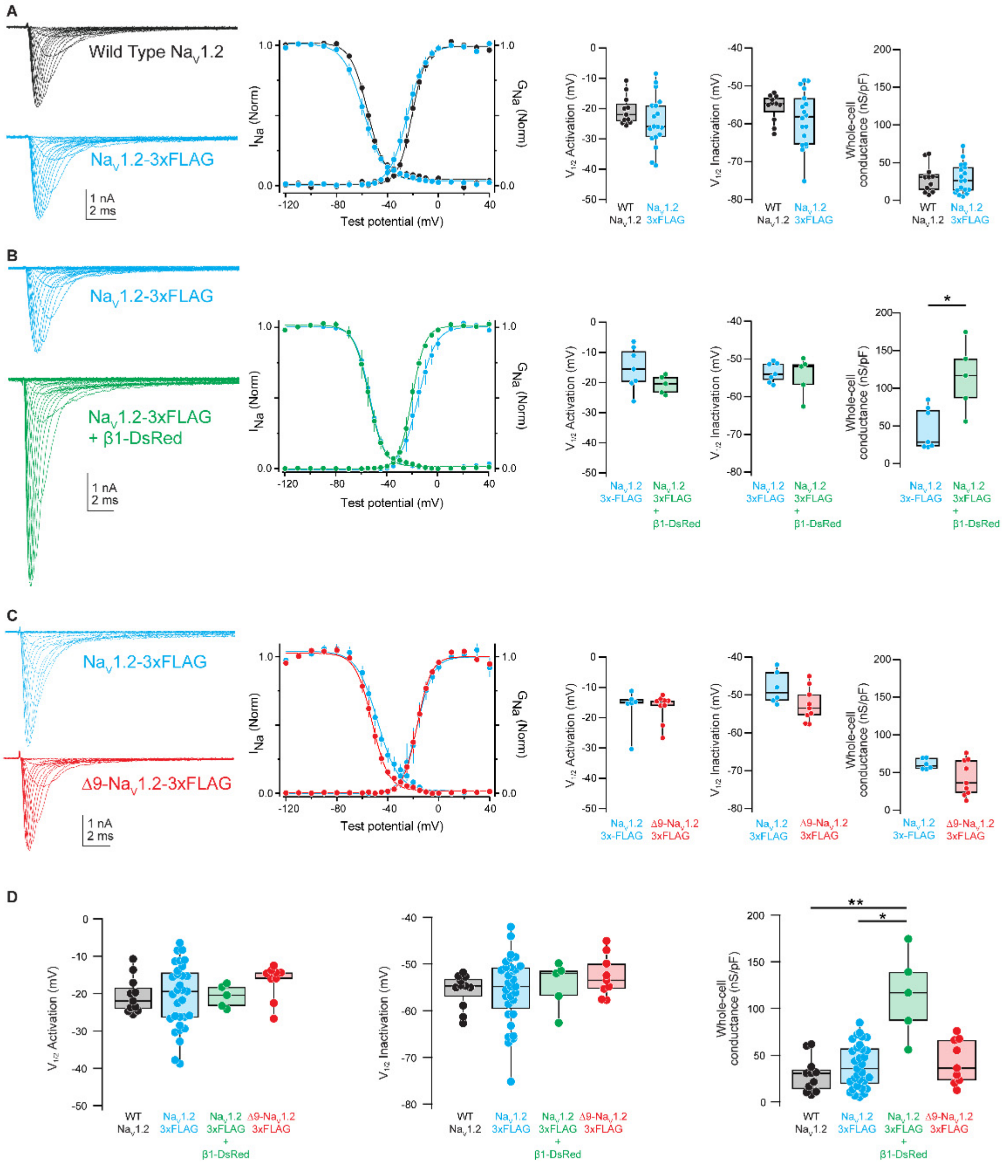


**Figure S1: Electrophysiological measurements of HEK293 cells transfected with $Na_V$1.2-IRES-eGFP constructs**

(A) Representative currents (left) and activation and inactivation curves (middle) from HEK293 cells transfected with WT $Na_V$1.2 (black) or $Na_V$1.2-3xFLAG-IRES-eGFP (blue). Right: Activation (left)- (WT $Na_V$1.2: -20.5 ± 1.5 mV, n = 11 cells; $Na_V$1.2-3xFLAG: -24.1 ± 2.0 mV, n = 18 cells). p = 0.15. Mann-Whitney test. Inactivation (middle)- (WT $Na_V$1.2: -55.7 ± 1.1 mV, n = 11; $Na_V$1.2-3xFLAG: -58.8 ± 1.7 mV, n = 18). p = 0.26. Mann-Whitney test. Whole cell conductance (right) – (WT $Na_V$1.2: 29.1 ± 5.6 nS/pF, *n* = 11; $Na_V$1.2-3xFLAG: 28.8 ± 4.6 nS/pF, n = 18). p = 0.95. Mann-Whitney test.

(B) Representative currents (left) and activation and inactivation curves (middle) from HEK293 cells transfected with $Na_V$1.2-3xFLAG-IRES-eGFP (blue) or $Na_V$1.2-3xFLAG-IRES-eGFP plus β1-DsRed (green). Right: Activation (left)- ($Na_V$1.2-3xFLAG: -15.3 ± 2.7 mV, n = 7 cells; $Na_V$1.2-3xFLAG + β1-DsRed: -20.7 ± 3.0 mV, n = 5 cells). p = 0.20. Mann-Whitney test. Inactivation (middle)- ($Na_V$1.2-3xFLAG: -53.5 ± 1.0 mV, n = 7; $Na_V$1.2-3xFLAG + β1-DsRed: -54.5 ± 2.3 mV, n = 5). p = 0.88. Mann-Whitney test. Whole cell conductance (right) – ($Na_V$1.2-3xFLAG: 46.2 ± 10.6 nS/pF, *n* = 7; $Na_V$1.2-3xFLAG + β1-DsRed: 114.7 ± 20.5 nS/pF, n = 5). * p = 0.018. Mann-Whitney test.

(C) Representative currents (left) and activation and inactivation curves (right) from HEK293 cells transfected with $Na_V$1.2-3xFLAG-IRES-eGFP (blue) or Δ9-$Na_V$1.2-3xFLAG-IRES-eGFP (red). Activation (left)- ($Na_V$1.2-3xFLAG: -16.8 ± 2.8 mV, n = 6 cells; Δ9-$Na_V$1.2-3xFLAG: -16.8 ± 1.6 mV, n = 9 cells). p = 0.95. Mann-Whitney test. Inactivation (middle)- ($Na_V$1.2-3xFLAG: -48.1 ± 1.7 mV, n = 6; Δ9-$Na_V$1.2-3xFLAG: -52.6 ± 1.5 mV, n = 9). p = 0.09. Mann-Whitney test. Whole cell conductance (right) – ($Na_V$1.2-3xFLAG: 39.5 ± 6.9 nS/pF, n = 6; Δ9-$Na_V$1.2-3xFLAG: 43.0 ± 7.9 nS/pF, n = 9). p = 0.65. Mann-Whitney test.

(D) Summary data from panels A-C. Activation (left)- (WT $Na_V$1.2: -20.5 ± 1.5 mV, n = 11 cells; $Na_V$1.2-3xFLAG: -20.7 ± 1.6 mV, n = 31 cells; $Na_V$1.2-3xFLAG + β1-DsRed: -20.7 ± 3.0 mV, n = 5 cells; Δ9-$Na_V$1.2-3xFLAG: -16.8 ± 1.6 mV, n = 9 cells). p = 0.46. Kruskal-Wallis test. Inactivation (middle)- (WT $Na_V$1.2: -55.7 ± 1.1 mV, n = 11; $Na_V$1.2-3xFLAG: -55.5 ± 1.3 mV, n = 31, $Na_V$1.2-3xFLAG + β1-DsRed: -54.5 ± 2.3 mV, n = 5; Δ9-$Na_V$1.2-3xFLAG: -52.6 ± 1.5 mV, n = 9). p =0.67. Kruskal-Wallis test. Whole cell conductance (right) – (WT Na$_V$1.2: 29.1 ± 5.6 nS/pF, *n* = 11; Na$_V$1.2-3xFLAG: 39.0 ± 4.2 nS/pF, n = 31; Na$_V$1.2-3xFLAG + β1-DsRed: 114.7 ± 20.5 nS/pF, n = 5; Δ9-Na$_V$1.2-3xFLAG: 43.0 ± 7.9 nS/pF, n = 9). p = 0.0065. Kruskal-Wallis test. Dunn's Multiple comparisons: * p=0.01 Na$_V$1.2-3xFLAG vs Na$_V$1.2-3xFLAG + β1-DsRed. ** p=0.004 Na$_V$1.2-IRES-GFP vs Na$_V$1.2-3xFLAG + β1-DsRed.

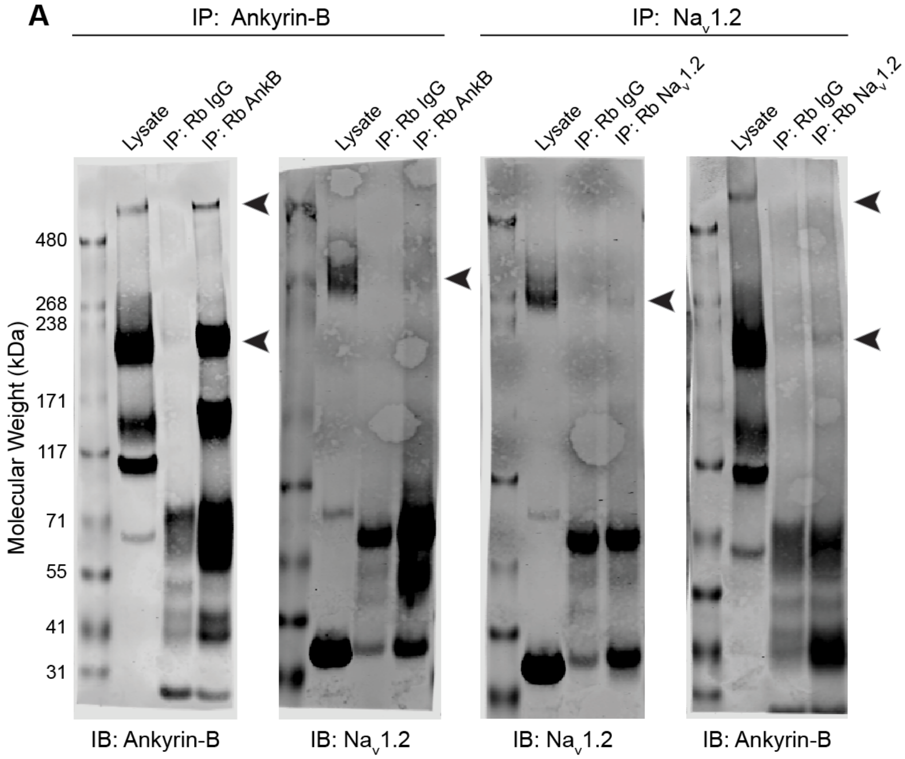


**Figure S2: Co-Immunoprecipitation of endogenous ankyrin-B and Na$_V$1.2 from adult mouse brain**

(A) Full membranes of Co-IP western blots from Figure 3E. Left: IP of endogenous ankyrin-B and western blot of IP lysates probed with antibodies to ankyrin-B or endogenous Na$_V$1.2 from P60-P75 mice. Right: IP of endogenous Na$_V$1.2 and western blot of IP lysates probed with antibodies to Na$_V$1.2 or ankyrin-B from P60-P75 mice. Black arrows highlight bands of ankyrin-B or Na$_V$1.2. Non-immune IgG used as a negative control.

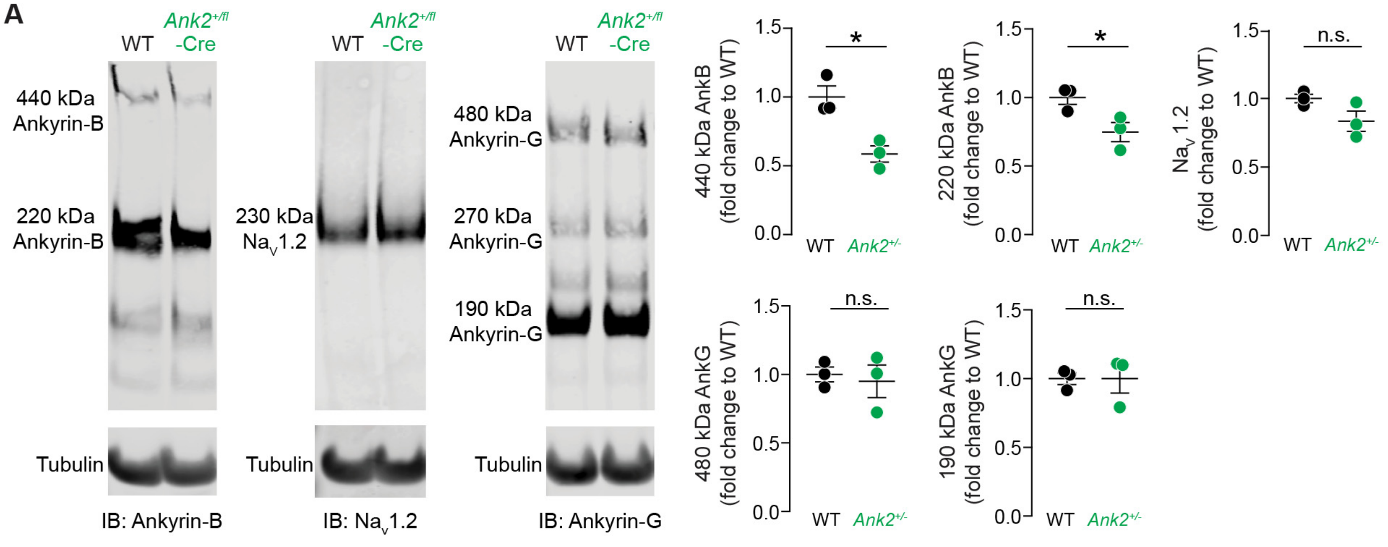


**Figure S3: Western blot of ankyrins and NaV1.2 from *Ank2*$^{+/fl}$::CaMKIIα-Cre adult mouse brain**

(A) Western blot of neocortical lysates of P60 WT (black) and *Ank2*$^{+/fl}$::CaMKIIα-Cre (green) mice. Left: Probed with anti-total ankyrin-B and anti-α-tubulin. 440 kDa ankyrin-B – (WT: 1.0 ± 0.08, N = 3 mice; *Ank2*$^{+/fl}$-Cre: 0.6 ± 0.06, N = 3 mice). p = 0.1. Mann-Whitney test. 220 kDa ankyrin-B – (WT: 1.0 ± 0.05, N = 3 mice; *Ank2*$^{+/fl}$-Cre: 0.75 ± 0.07, N = 3 mice). p = 0.1. Mann-Whitney test. Middle: Probed with anti-NaV1.2 and anti-tubulin. NaV1.2 – (WT: 1.0 ± 0.03, N = 3 mice; *Ank2*$^{+/fl}$-Cre: 0.83 ± 0.07, N = 3 mice). p = 0.2. Mann-Whitney test. Right: probed with anti-total ankyrin-G and anti-tubulin. 480 kDa ankyrin-B – (WT: 1.0 ± 0.05, N = 3 mice; *Ank2*$^{+/fl}$-Cre: 0.95 ± 0.1, N = 3 mice). p > 0.99. Mann-Whitney test. 190 kDa ankyrin-B – (WT: 1.0 ± 0.04, N = 3 mice; *Ank2*$^{+/fl}$-Cre: 1.0 ± 0.1, N = 3 mice). p = 0.7. Mann-Whitney test.

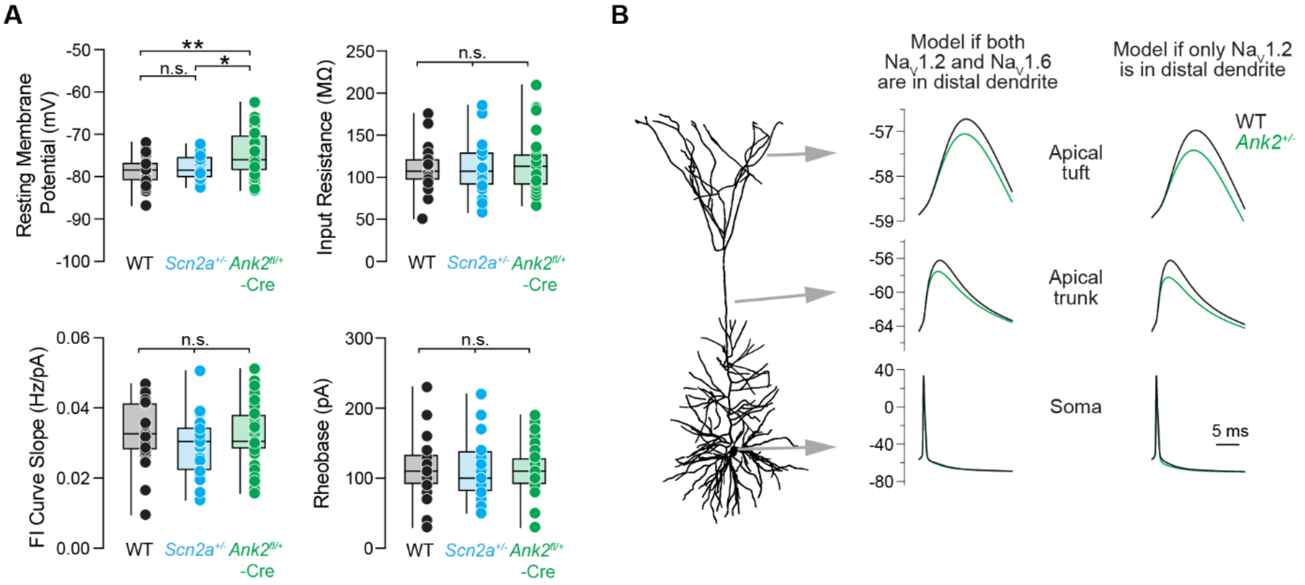


**Figure S4: Additional modeling and intrinsic electrophysiological measurements from conditional *Ank2*$^{+/fl}$-Cre mice**

(A) Empirical electrophysiological recordings from P43-75 WT (black), *Scn2a*$^{+/-}$ (cyan) vs. *Ank2*$^{+/fl}$-Cre (green) L5 thick-tufted pyramidal neurons. Circles represent single cells. Resting membrane potential (mV) – (WT: -78.7 ± 0.6 mV, n = 27 cells; *Scn2a*$^{+/-}$: -77.7 ± 0.6 mV, n = 23 cells; *Ank2*$^{+/fl}$-Cre: -74.9 ± 0.8 mV, n = 39 cells). WT vs. *Scn2a*$^{+/-}$ p = 0.7, WT vs *Ank2*$^{+/fl}$-Cre **p = 0.001, *Scn2a*$^{+/-}$ vs. *Ank2*$^{+/fl}$-Cre *p = 0.03. Holm-Šídák multiple comparisons test. Input resistance (MΩ) – (WT: 109.9 ± 5.0 MΩ, n = 27 cells; *Scn2a*$^{+/-}$: 109.9 ± 5.7 MΩ, n = 27 cells; *Ank2*$^{+/fl}$-Cre: 114.5 ± 5.3 MΩ, n = 39 cells). No significant differences. Holm-Šídák multiple comparisons test. FI curve slope (Hz/pA) - (WT: 0.033 ± 0.002 Hz/pA, n = 29 cells; *Scn2a*$^{+/-}$: 0.03 ± 0.002 Hz/pA, n = 23 cells, *Ank2*$^{+/fl}$-Cre: 0.033 ± 0.001, n = 39 cells). No significant differences. Holm-Šídák multiple comparisons test. Rheobase rheobase current (pA) to generate first spike – (WT: 116.6 ± 7.9 pA, n = 29 cells; *Scn2a*$^{+/-}$: 115.2 ± 10.0 pA, n = 23 cells; *Ank2*$^{+/fl}$-Cre: 112.1 ± 5.6 pA, n = 39 cells). No significant differences. Holm-Šídák multiple comparisons test.

(B) Compartmental modeling of the effects of *Ank2* haploinsufficiency, and presumed resultant 50% reduction in $Na_V1.2$ density in regions of dendrite >30 µm from the soma. Two models are considered: a model similar to that in Spratt et al., 2019 where both $Na_V1.2$ and $Na_V1.6$ are expressed in equal densities in dendrites (left), and a revised model in which $Na_V1.2$ is the only sodium channel proteoforms expressed in distal dendrites. In both cases, *Ank2*$^{+/-}$ conditions are modeled as a 50% reduction in $Na_V1.2$ density within more distal dendritic compartments only (e.g., soma and proximal dendrites maintain WT levels of $Na_V1.2$).

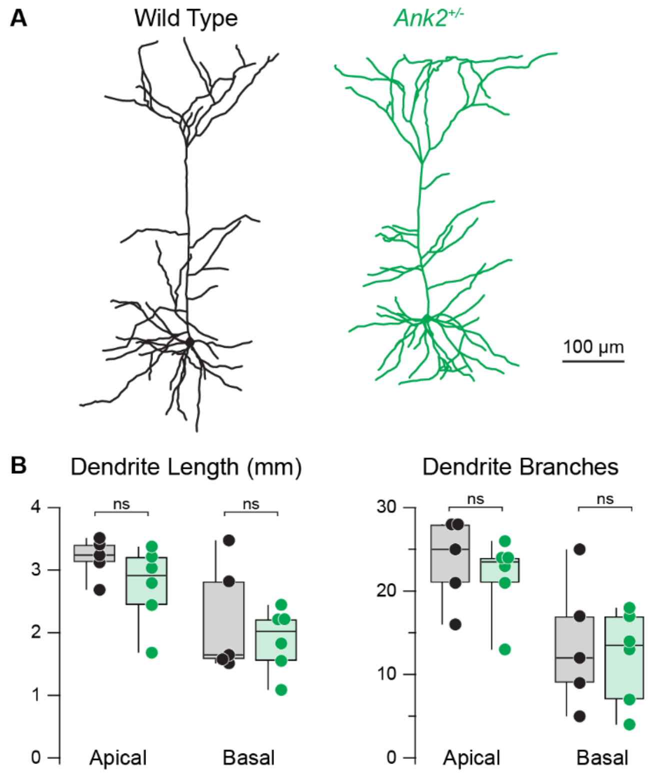


**Figure S5: Dendritic morphology and arborization of *Ank2*$^{+/fl}$-Cre neurons**

(A) Example tracings of L5 thick-tufted neurons from P52-67 WT (black) and *Ank2*$^{+/fl}$-Cre (green) mice generated from 2PLSM z-stacks of Alexa 594-filled neurons.

(B) Left: Dendritic length (mm) of apical and basal dendrites. Apical – (WT: 3.2 ± 0.1 mm, n = 5 cells; *Ank2*$^{+/fl}$-Cre: 2.8 ± 2.5 mm, n = 6 cells). p = 0.18. Mann-Whitney test. Basal – (WT: 2.2 ± 0.4 mm, n = 5 cells; *Ank2*$^{+/fl}$-Cre: 1.9 ± 0.2 mm, n = 6 cells). p = 0.8. Mann-Whitney test. Right: Overall number of branch points per cell. Apical – (WT: 23.6 ± 2.3, n = 5 cells; *Ank2*$^{+/fl}$-Cre: 21.8 ± 1.9, n = 6 cells). p = 0.46. Mann-Whitney test. Basal – (WT: 13.6 ± 3.5, n = 5 cells; *Ank2*$^{+/fl}$-Cre: 12.17 ± 2.3, n = 6 cells). p = 0.97. Mann-Whitney test.

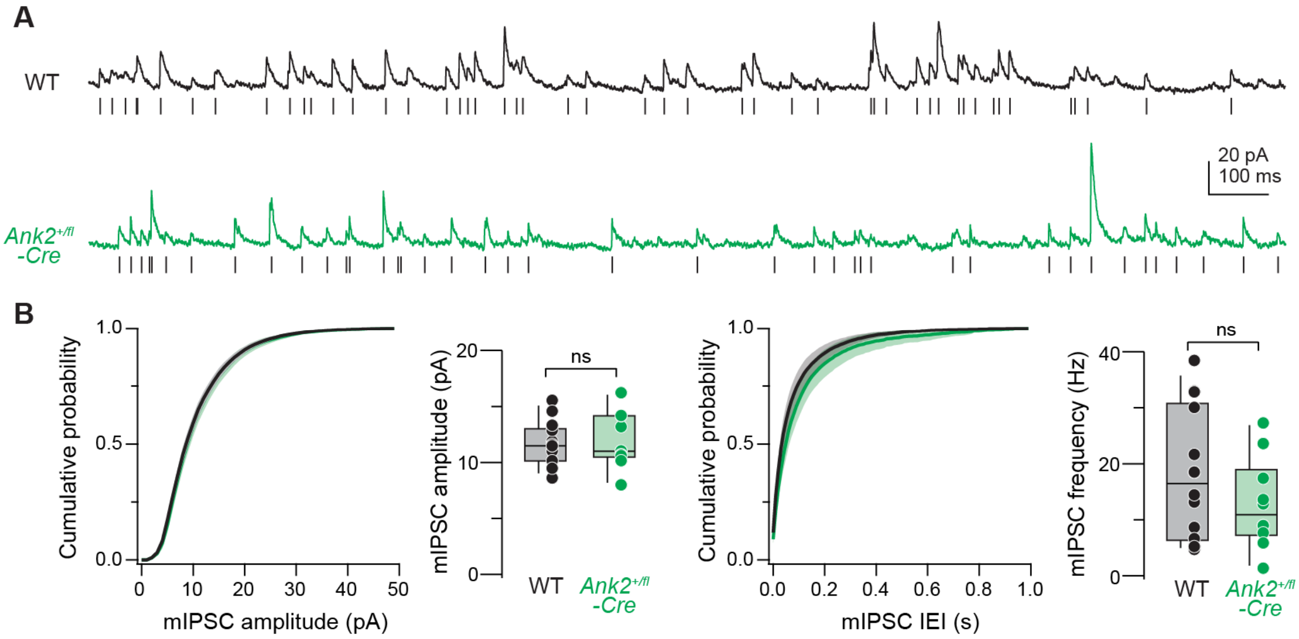


**Figure S6: Miniature inhibitory postsynaptic currents from *Ank2*$^{+/fl}$-Cre L5 pyramidal neurons.**

(A) mIPSCs recorded from P54-P60 WT (black) and *Ank2*$^{+/fl}$-Cre-mCherry-positive (green) L5 pyramidal neurons at 0 mV. Ticks denote detected events. Cumulative probability histograms were generated per cell and then averaged. Circles represent individual neurons. Left: Cumulative probability distribution and mIPSC amplitude (pA) – (WT: 11.7 ± 0.5 pA, n = 14 cells; *Ank2*$^{+/fl}$-Cre: 11.9 ± 0.8 pA, n = 10 cells). p > 0.99. Mann-Whitney test. Right: Cumulative probability distribution of mIPSC inter-event intervals (IEI) and average frequency (Hz) – (WT: 17.9 ± 3.1 Hz, n = 14 cells; *Ank2*$^{+/fl}$-Cre: 12.8 ± 2.5 Hz, n = 10 cells). p = 0.4. Mann-Whitney test.
